# Proteome-wide non-cleavable crosslink identification with MS Annika 3.0 reveals the structure of the *C. elegans* Box C/D complex

**DOI:** 10.1101/2024.09.03.610962

**Authors:** Micha J. Birklbauer, Fränze Müller, Sowmya Sivakumar Geetha, Manuel Matzinger, Karl Mechtler, Viktoria Dorfer

**Author notes:** Contributing authors.

## Abstract

The field of crosslinking mass spectrometry has seen substantial advancements over the past decades, enabling the structural analysis of proteins and protein-complexes and serving as a powerful tool in protein-protein interaction studies. However, data analysis of large non-cleavable crosslink studies is still a mostly unsolved problem due to its *n*-squared complexity. We here introduce a novel algorithm for the identification of non-cleavable crosslinks implemented in our crosslinking search engine MS Annika that is based on sparse matrix multiplication and allows for proteome-wide searches on commodity hardware. Application of this new algorithm enabled us to employ a proteome-wide search of *C. elegans* nuclei samples, where we were able to uncover previously unknown protein interactions and conclude a comprehensive structural analysis that provides a detailed view of the Box C/D complex, enhancing our understanding of its assembly and functional dynamics. Our findings provide valuable insights into the intricate regulation of cellular homeostasis and immune responses, which are conserved across species, including humans. Moreover, our algorithm will enable researchers to conduct similar studies that were previously unfeasible.

## 1 Introduction

The past decade has seen continuous improvements in the field of crosslinking mass spectrometry (XLMS) on both the experimental [1, 2] and the data analysis side [3, 4] with numerous new software being released for crosslink identification [5–7]. Today the technique of XLMS has matured into a powerful tool for structural, molecular, and systems biology [8] which enables the structural analysis of proteins and proteincomplexes [9, 10], as well as capturing protein-protein interactions potentially up to system-wide scale, allowing in-depth studies of large interactomes [11, 12]. Plenty of comprehensive reviews highlighting successful XLMS applications, potential pitfalls and drawbacks have been published in recent years [13–18]. However, while XLMS has seen substantial advancements, there is one major challenge that still exists: computational analysis of mass spectra originating from non-cleavable crosslinking reagents poses a complex problem that is largely unexplored due to its *n*-squared nature. Contrary to cleavable crosslinkers like DSSO [19] and DSBU [20], non-cleavable reagents such as DSS [21] and BS3 [22] do not incorporate an off-centre labile moiety that is cleaved during fragmentation and as a result do not yield characteristic doublet ions required for mass calculation of the individual crosslinked peptides. Therefore, only the mass of the complete non-cleavable crosslinked entity is known and all peptide combinations arising from the protein database that match the entity’s mass have to be considered for identification. The number of combinations grows with the square of the protein database size, hence the name *n*-squared problem [23]. Moreover, accurate estimation of the false discovery rate (FDR) used for crosslink validation is also more complex for non-cleavable crosslinks as peptide identifications are interdependent on each other [24]. This behaviour renders large proteome-wide studies with non-cleavable crosslinkers largely unfeasible, ultimately because most crosslink search engines are unable to deal with the enormous search spaces that have to be accounted for in such studies. Nevertheless, despite the computational challenges, non-cleavable crosslinkers are still the most used class of crosslinking reagents [25] due to very distinct advantages: non-cleavable crosslinkers are simpler in chemical structure and hence easier to synthesise, making them the more cost-effective option compared to cleavable reagents. For large-scale studies or routine analyses, non-cleavable crosslinkers offer a budget-friendly option without compromising the quality of the data [15]. Furthermore, because of their simpler structure, non-cleavable crosslinkers also feature less chemical groups that are prone to introducing potential side reactions which are undesirable especially in whole cell crosslinking experiments. Contrary to cleavable crosslinkers, non-cleavable reagents do not require optimization of mass spectrometry worflows for detection of signature ions that are integral for cleavable searches [26]. Most importantly, a non-cleavable crosslinker’s backbone is stable and chemically inert to side reactions during sample preparation, making them suitable for applications where structural integrity and long-term stability are critical. Their usage is also in many cases attractive because of their chemical properties such as various levels of hydrophobicity and membrane permeability. This highlights the need for an efficient and robust crosslinking search engine capable of analysing data from complex non-cleavable crosslink samples such as proteome-wide studies.

One such biological system that would greatly benefit of the insights from non-cleavable crosslinking is the *Caenorhabditis elegans* nucleus, specifically the Box C/D ribonucleoprotein (RNP) complex which is a crucial molecular assembly involved in RNA modification processes, primarily catalysing the 2’-O-methylation of specific nucleotides in ribosomal RNA (rRNA). This modification is essential for the proper functioning of ribosomes, the cellular machinery responsible for protein synthesis. The complex is composed of small nucleolar RNAs (snoRNAs) and core proteins, including fib-1 (fibrillarin), nol-56 (Nop56), nol-58 (Nop58), and SNU13 (M28.5) [27]. These components assemble into a functional unit that directs the methylation machinery to precise sites on the pre-rRNA, guided by sequence complementarity between the snoRNA and the target rRNA sequence [28]. Recent studies have uncovered a novel role for the Box C/D RNP complex beyond its traditional function in rRNA modification [29–32]. In *C. elegans*, this complex plays a significant role in mitochondrial surveillance and regulating innate immune responses [33]. Tjahjono *et al.* [33] have shown that Box C/D small nucleolar ribonucleoproteins (snoRNPs) are essential for the proper activation of mitochondrial stress response pathways, including the unfolded protein response in mitochondria (UPR*^mt^*) and the Ethanol and Stress Response Element (ESRE) network. These pathways are crucial for maintaining mitochondrial health and functionality, particularly under stress conditions. Understanding the multifaceted roles of the Box C/D RNP complex in *C. elegans* provides valuable insights into the intricate regulation of cellular homeostasis and immune responses, which are conserved across species, including humans [34]. This knowledge has potential implications for understanding the mechanisms underlying mitochondrial health, immune responses, and their impact on ageing and disease.

In the following study we present a new algorithm, MS Annika 3.0, for proteome-wide identification of non-cleavable crosslinks which we employ to study the structural arrangement and interaction landscape of the *C. elgeans* Box C/D RNP complex, uncovering new insights.

## 2 Results

We here introduce a novel algorithm for identification of non-cleavable crosslinks implemented in our search engine MS Annika [5, 35] which is capable of analysing proteome-wide studies in reasonable time on commodity hardware. Moreover, to show the applicability of this approach we performed crosslinking experiments with *C. elegans* nuclei and successfully searched the mass spectrometry data against the full *C. elegans* proteome of over 26 000 proteins. Using the identified crosslinks we were able to conclude a comprehensive structural analysis that provides a detailed view of the Box C/D RNP complex in *C. elegans*, enhancing our understanding of its assembly and functional dynamics. Finally, we compared our new algorithm against other state-of-the-art crosslinking search engines commonly used in the field and proved that MS Annika is either on par or better than competing tools.

### 2.1 Implementation of a novel algorithm for identification of non-cleavable crosslinks

Identification of crosslinks originating from non-cleavable crosslinkers poses a big computational challenge due to the combinatorial explosion of the search space that we describe earlier. In order to successfully apply crosslink identification beyond human proteome-wide scale, crosslink search engines need to be extremely efficient in their choice of peptide candidates to consider for crosslink detection. We here introduce a novel algorithm implemented in our search engine MS Annika that can accurately determine good peptide candidates for crosslink identification in reasonable time on commodity hardware, enabling crosslink searches for protein databases of up to 20 million peptides. Figure 1 shows a schematic overview of the complete algorithm as implemented in MS Annika. At the core, this algorithm approximately scores all peptides in the database against every experimental mass spectrum and yields the top candidates for each spectrum to consider for crosslink search which drastically reduces the search space. This approximate scoring of all peptides against a spectrum is possible because peptides and spectra are represented as (sparse) vectors and the problem of calculating the scores of every peptide in the database can be denoted as a simple sparse matrix multiplication. An in-depth description of the algorithm is given in Section 4.1.

**Fig. 1.**
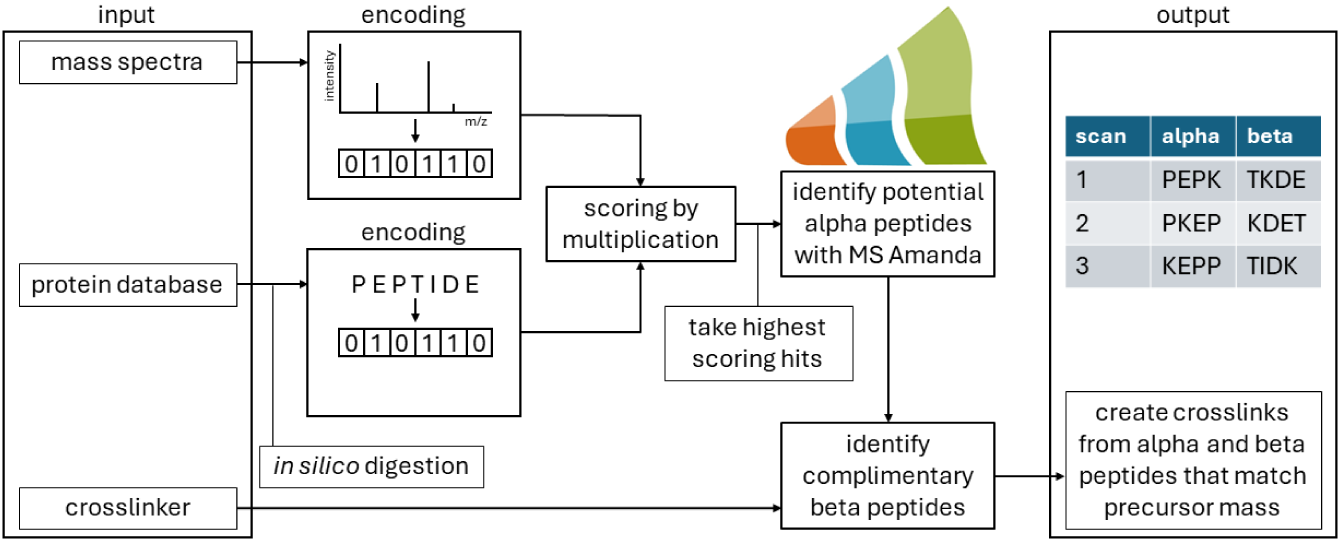
Schematic overview of the algorithm for identification of non-cleavable crosslinks in our search engine MS Annika [5, 35]. Mass spectra and peptides arising from the *in silico* digestion of the protein database are encoded as sparse vectors and subsequently scored by matrix multiplication. The highest scoring hits are considered for the identification of potential alpha peptides with our inhouse developed peptide search engine MS Amanda [36, 37]. Identified alpha peptides are used to find complimentary beta peptides and ultimately alpha and beta peptides matching the mass spectrum’s precursor mass are combined to crosslinks.

### 2.2 Identification of crosslinks in *C. elegans* using a proteome-wide non-cleavable search

The main advantage of the new non-cleavable search algorithm in MS Annika is its ability to efficiently process very large protein databases. This is an almost exclusive feature of MS Annika since most non-cleavable crosslink search engines are not able to handle protein databases consisting of more than a couple of thousand proteins. In order to demonstrate the applicability of MS Annika for large proteome-wide studies, we searched mass spectrometry data of *C. elegans* nuclei that we crosslinked with DSG against the full *C. elegans* proteome, amounting to 26 695 proteins in total. Furthermore, to study the impact of protein database size we compared the results against the same search using a filtered proteome containing only proteins that occurred in more than two high-confidence peptide-spectrum-matches (PSMs) in a preliminary linear search for identification of non-crosslinked peptides. The filtered proteome consisted of 3069 proteins in total. Figure 2 shows the number of identified crosslinks at 1% estimated FDR per biological replicate for both protein databases using MS Annika for identification and xiFDR [24] for validation: remarkably, despite the large difference in protein database size, the number of identified crosslinks at 1% estimated FDR shows only small variation regardless of the used database. The biggest change is observed for replicate three where using the filtered proteome causes a gain of 39 crosslinks, resulting in 249 crosslinks instead of 210. The total number of identified unique crosslinks at 1% estimated FDR across all three biological replicates is 448 when using the filtered proteome and 435 when using the full proteome. Supplementary Figure 1 shows the overlaps in crosslink identifications between the replicates and between filtered and full proteome identifications with generally good agreement.

**Fig. 2.**
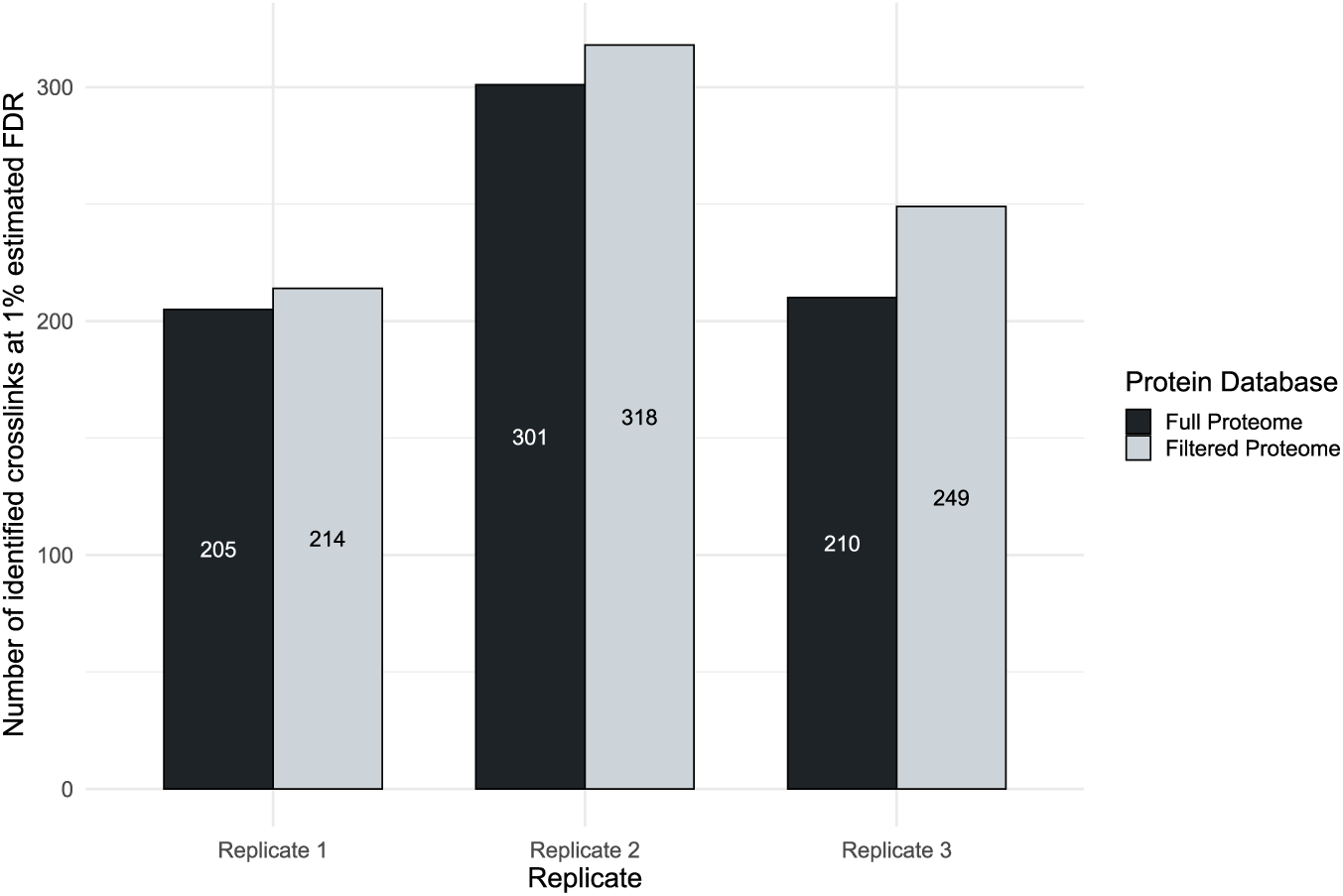
Influence of protein database size on the non-cleavable search of MS Annika. Mass spectrometry data of *C. elegans* nuclei was searched with MS Annika once using the full *C. Elegans* proteome (n = 26 695) and once using a filtered proteome of abundant proteins identified in a linear search (n = 3069). Results were validated for 1% estimated FDR with xiFDR [24]. Interestingly, the number of identified crosslinks does not show large variation, regardless of database size with the biggest difference observed in replicate three where the filtered database search reports 39 more crosslinks. Overall 448 unique crosslinks are identified across the three replicates using the filtered proteome and 435 unique crosslinks using the full proteome which shows the applicability of MS Annika for large proteome-wide studies.

Supplementary Figure 2 and 3 depict the results using MS Annika with the built-in validation algorithm instead of validation with xiFDR: the differences in results between using the filtered and full proteome are more pronounced, especially for replicate one where using the full proteome for search causes a loss of more than half of the identified crosslinks. Initially it might seem counter-intuitive that usage of the larger database yields less crosslinks, however as the size of the database grows, so does the chance of randomly matching a false positive hit which has to be accounted for during validation. Specifically, the greater chance to match false positive hits results in higher score cut-offs to preserve the 1% FDR threshold and therefore leads to less reported crosslinks. This highlights the benefit of using more sophisticated validation tools like xiFDR for larger protein databases which we explore further in Section 2.6.

All searches were run on a desktop PC with moderate hardware (12-core AMD Ryzen R9 7900X 4.7GHz CPU with 64 GB of memory) at a mean runtime of 6 hours and an average of 549 460 mass spectra per replicate for the full proteome-wide search. The exact time measurements for each replicate and hardware specifications are given in Supplementary Table 1 and 2, respectively. It should be noted however, that even though the system had 64 GB of memory installed, the full memory capacity was never utilized. In fact, such a proteome-wide search does run on a normal laptop with a 4-core CPU and 16 GB of memory, albeit MS Annika does heavily benefit of additional CPU cores due to the parallel nature of the algorithm.

### 2.3 Structural analysis of the Box C/D RNP complex in *C. elegans*

The structural organisation and interaction network of the Box C/D RNP complex in *C. elegans* were elucidated through an integrative approach combining XLMS using the data and results as described in Section 2.2 and structural modelling. The interaction map, presented in Figure 3, highlights the spatial arrangement and connectivity among the core components of the complex, including nol-56, nol-58, and fib-1, along with M28.5 and NEDG-01330. Figure 3A illustrates an AlphaFold2 Multimer [38–41] screening of potential complex interactors with nol-58 as bait protein against a fasta file containing 29 known and potential interactors of the Box C/D RNP complex. The screening results are shown with all nol-58 candidate predictions (n = 29) as circles and ranked by the average interface prediction TM score (interface pTM). Interestingly, the prediction of the core complex revealed two new potential interactors NEDG-01330 (top hit) and NEDG-01670 (fourth top hit). Additionally, nol-58 establishes robust interactions with both nol-56 and fib-1 as well as M28.5, consistent with its central role in the complex assembly known in the literature (Figure 3E) [33]. The interaction of the core complex was further confirmed by crosslinking mass spectrometry as shown in Figure 3B. Identified crosslink sites were mapped onto their respective protein sequences using xiView [42]. While nol-58 exhibits only one protein-protein interaction (PPI) link to nol-56, it established multiple interaction sites with fib-1, reinforcing their cooperative function within the complex. The snoRNP M28.5, with its distinct secondary structure, anchors these proteins, facilitating the formation of a stable and functional RNP complex. Crosslink restrains could not be identified by mass spectrometry for NEDG-01330 and NEDG-01670 although the AlphaFold2 Multimer screening predicted both proteins as strong interactors. Remarkably, the predicted tertiary structure of NEDG-01330 is very similar to M28.5 and its predicted position in the complex suggests a similar role in complex assembly compared to M28.5 (Figure 3C). The fib-1 protein, represented in light pink, occupies a central position, interacting with both nol-56 (firebrick) and nol-58 (orange), while M28.5 (pink) wraps around these proteins, ensuring their proper orientation and function within the RNP assembly. The predicted three-dimensional structural model of the Box C/D RNP complex, integrating the crosslinking data, shows a clear violation of PPI links (light green) exceeding the maximum allowed distance of 19-22 Å, resulting from the DSG crosslinker backbone of 7.7 Å and two times the lysine side chain of 6-7 Å (Figure 3C). Hence, to refine the structure of the Box C/D RNP complex we employed AlphaLink2 [43, 44] and integrated our crosslink restraints into the prediction process. This resulted in a refined model with crosslink restraints fulfilling the distance limit of 22 Å except for two inter-crosslinks that remained violated (Figure 3D).The refinement by AlphaLink2 demonstrates a significant improvement in structure prediction, as indicated by an increase in the ipTM score from 0.689 to 0.721, accompanied by a reduction in crosslink violations (from 3 to 2) and, more importantly, shorter distances for all crosslinks (Figure 3F). Despite the refinement, two crosslinks still exceeded the distance limit due to the inherent symmetry of the complex. As illustrated in Figure 3E, the complex consists of two M28.5 and two fib1 proteins, one on each side. The long-distance crosslink between M28.5 and nol-56 can be attributed to a second M28.5 protein interacting with the C-terminal region of nol-56, like the predicted interaction between M28.5 and nol-58. This interaction with a second M28.5 protein would satisfy the crosslink distance limit. However, due to limitations in the number of complex members that can be provided for structure prediction, it was not possible to test this hypothesis. The same symmetrical consideration applies to the long-distance link between fib-1 and nol-58, with the potential formation of a new interaction interface between fib-1 and NEDG-01330. It is plausible that fib-1 and NEDG-01330 form a dimer that binds to nol-58 at residue 194 and adjacent residues. Although the predicted model of the Box C/D snoRNP complex raises questions that require further investigation in future experiments, the structural analysis offers a detailed and improved view of the Box C/D RNP complex in *C. elegans* and significantly enhances our understanding of the assembly and functional dynamics of the complex.

**Fig. 3.**
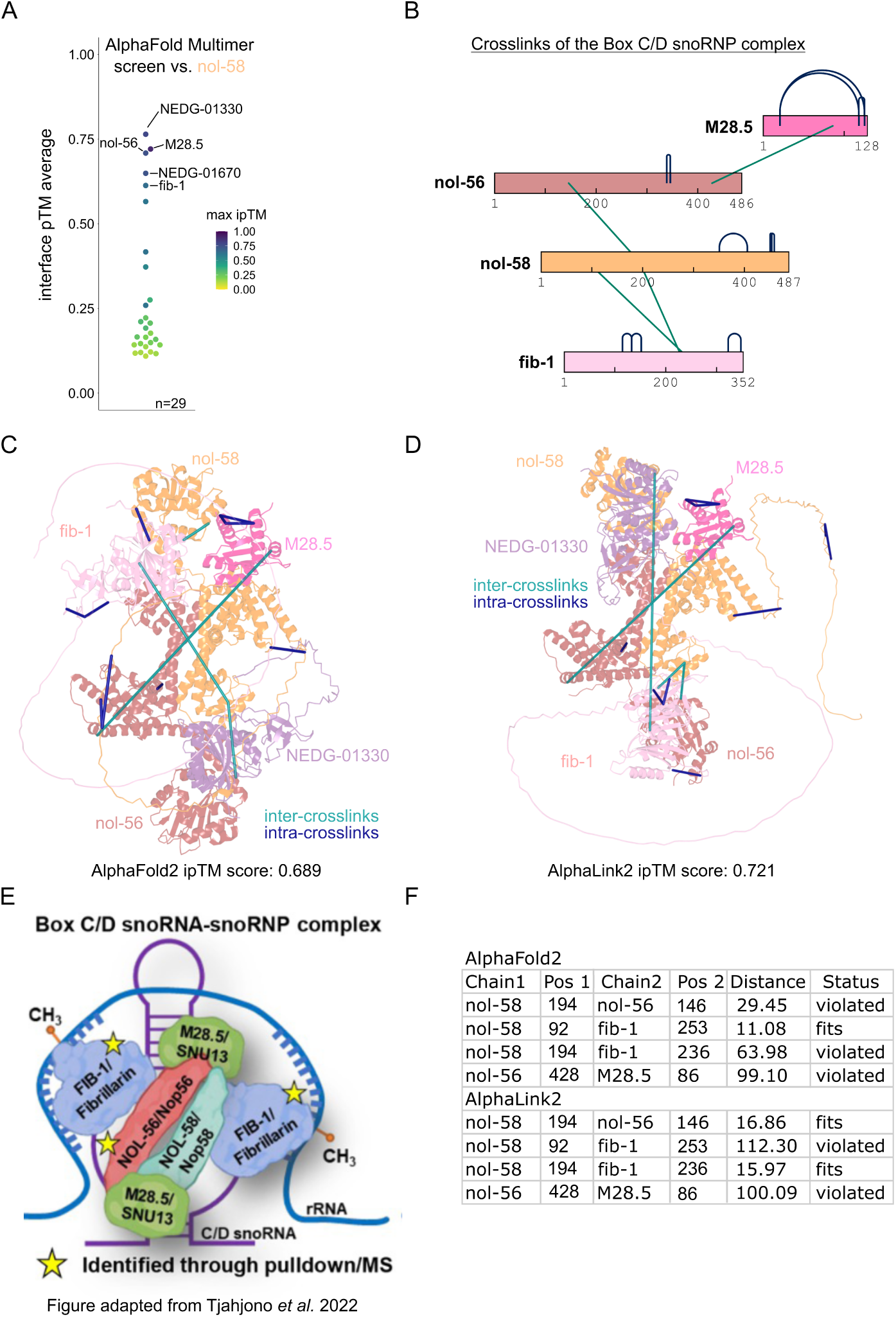
Modelling and structure refinement of the Box C/D RNP complex using AlphaFold2 and AlphaLink2. (A) AlphaFold2 Multimer [38–41] protein interaction screen to identify and confirm interactors of nol-58. The screening results are shown with all nol-58 candidate predictions (n = 29) as circles and ranked by the average interface prediction TM score (interface pTM). The colour code represents the best ipTM score across all five structure predictions per protein. The top five predicted interactors are highlighted with corresponding gene names. (B) Crosslink visualisation by xiView [42] of identified links from the Box C/D RNP complex. Intra-protein links are coloured in dark blue while interlinks between proteins are coloured in light green. (C) AlphaFold2 prediction of the Box C/D RNP complex as cartoon schematic with nol-583i3n orange, M28.5 in pink, NEDG-01330 in violet, nol-56 in firebrick and fib-1 in light pink. Inter (light green) and intra (dark blue) crosslinks are plotted on top of the structure to visualise the interaction relations between the complex members. (D) AlphaLink2 refinement of the predicted Box C/D RNP complex with nol-58 in orange, M28.5 in pink, NEDG-01330 in violet, nol-56 in firebrick and fib-1 in light pink. Inter (light green) and intra (dark blue) crosslinks are plotted on top of the structure. (E) Figure adapted from Tjahjono *et al.* 2022 [33] showing the current Box C/D RNP complex model currently available in the literature. (F) Summary of inter-links after AlphaFold2 and AlphaLink2 prediction.

### 2.4 MS Annika accurately identifies non-cleavable crosslinks in benchmark datasets

We further evaluated the non-cleavable search algorithm of MS Annika by analysing the crosslinking benchmark datasets of Beveridge and co-workers [45] and Matzinger and co-workers [46] that were specifically designed to assess the quality of crosslinking search engines and which allow for the computation of an experimentally validated FDR. Moreover, we compared the results of MS Annika against other state-of-theart tools commonly used in the field for identification of non-cleavable crosslinks, namely MaxLynx [6] (part of MaxQuant [47]), MeroX [48, 49], pLink [50], xiSearch [51] (including xiFDR [24]), and XlinkX [52].

The dataset of Beveridge and co-workers [45] consists of three technical replicates of synthetic peptides from *Streptococcus pyogenes* Cas9 crosslinked with the crosslinker DSS [21]. The mass spectrometry data was searched against a protein database of Cas9 and 10 contaminant proteins. Figure 4 shows the results of the different tools at 1% estimated FDR: Reporting 218 unique true positive crosslinks MS Annika detects 8 less true positive crosslinks but also 7 less false positive crosslinks than MaxLynx on average across the three replicates, therefore - at 1.37% - yielding a 2.94% better experimentally validated FDR than MaxLynx. In contrast, while MeroX yields the best experimentally validated FDR at zero false positive hits, it also identifies only 46 unique true positive crosslinks on average. The crosslinking search engine pLink reports the highest number of true positive crosslinks on average at 242, however this is at the cost of 17 false positive hits and yielding an experimentally validated FDR of 6.52% on average which is the second worst only after XlinkX. Furthermore, at 220 true positive identifications, which is 2 more than MS Annika, and only 2 false positive crosslinks (1 less than MS Annika) on average xiSearch reports arguably the best result for this dataset, yielding an average experimentally validated FDR of 1.05%, very close to the target of 1% estimated FDR. On the other hand XlinkX returns what is possibly the worst result for all the compared tools with an average of 31 false positive identifications and an experimentally validated FDR of 15.19% while reporting 173 unique true positive crosslinks. Despite this being a rather simple dataset with only 11 proteins, MaxLynx, pLink and XlinkX noticeably underestimate the actual FDR, reporting a lot more false positives than allowed. Supplementary Figure 4 shows the intersection and union of results from MS Annika, MaxLynx and pLink for all three replicates including calculated experimentally validated FDRs. The venn diagrams show high agreement in identifications and intersections consist of zero false positive hits.

**Fig. 4.**
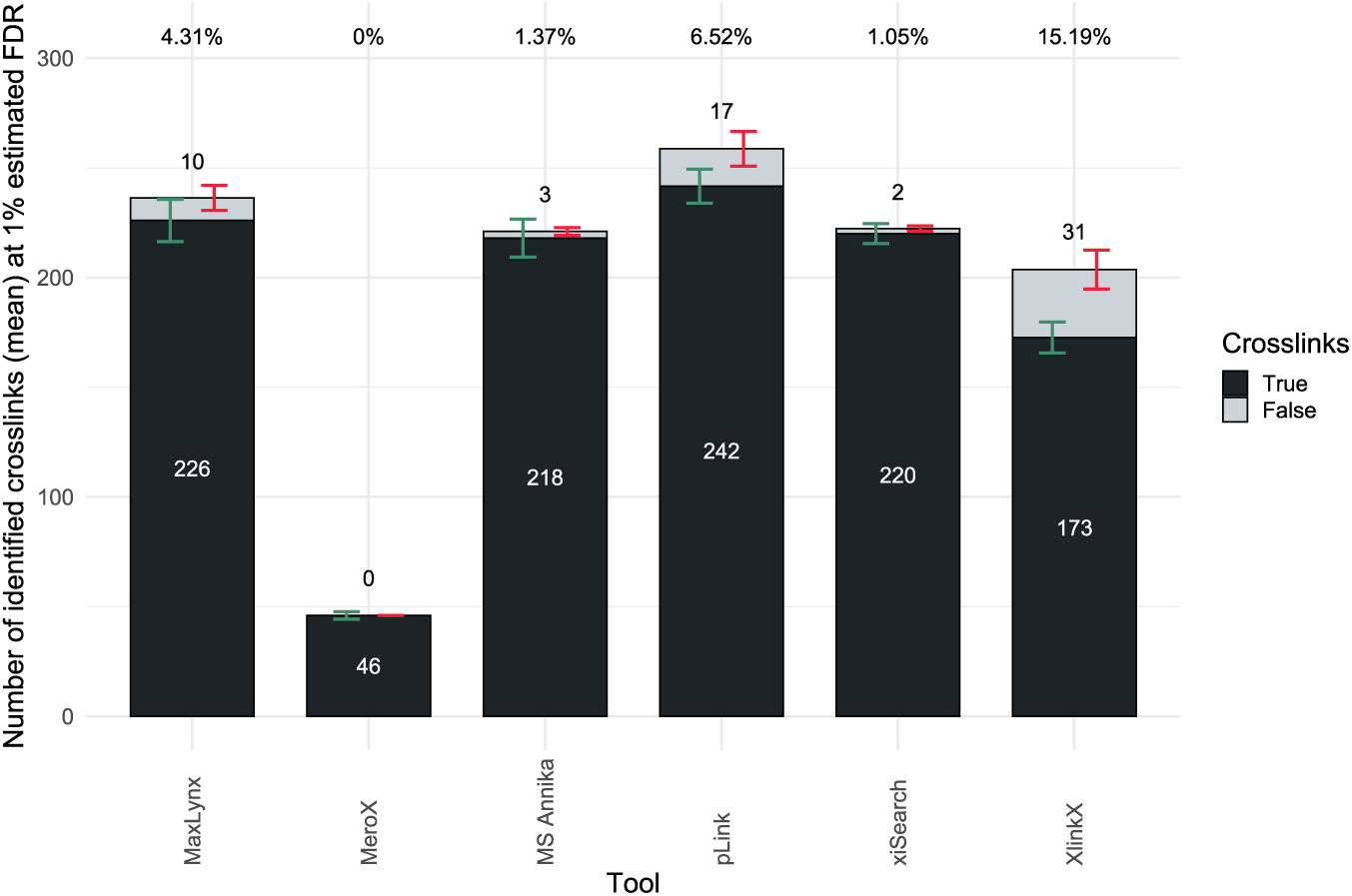
Identified crosslinks and experimentally validated FDRs of the different crosslinking search engines using the benchmark dataset by Beveridge and co-workers where synthetic peptides were crosslinked with DSS [45]. Both MS Annika and xiSearch [51] report very good results at around 220 true positive crosslink identifications and experimentally validated FDRs that are very close to the target FDR of 1%. On the other hand, both MaxLynx [6] and pLink [50] report more true positive crosslinks but at the cost of more false positive hits and higher experimentally validated FDRs of up to 6.52%, already far off the 1% target. XlinkX and MeroX are both outliers with very distinct results, XlinkX [52] yields a very high experimentally validated FDR of 15.19% while MeroX [48, 49] reports only 46 true positive crosslinks. Results were validated for 1% estimated FDR. All numbers are averages from three technical replicates (n = 3), the number of crosslinks was rounded to the closest integer value. Percentage numbers above the bars denote the average calculated experimentally validated FDR rounded to two decimal places. Error bars denote the standard deviation, in green for true positive crosslinks and in red for false positive crosslinks

The dataset by Matzinger and co-workers is more complex, attempting to more closely resemble real crosslinking experiments while still relying on synthetic peptides and therefore also allowing the calculation of an experimentally validatable FDR [46]. For this dataset synthetic ribosomal peptides from *Escherichia coli* were crosslinked with ADH, an acidic crosslinker primarily reacting with aspartic acid and glutamic acid. The mass spectrometry data consisting of three technical replicates was searched against 171 sequences of the *E. coli* ribosomal complex which is a database size that is large enough to pose a challenge for crosslinking search engines such as MaxLynx that use an exhaustive search for crosslink identification. Figure 5 shows the results achieved by the different crosslinking search engines on this dataset when validating for 1% estimated FDR. Evidently, all search engines are underestimating the actual FDR, reporting more false positive hits than what would be allowed at 1% FDR. For this dataset MS Annika yields the lowest experimentally validated FDR at a mean of 3.92%, identifying on average 89 unique true positive crosslinks and 4 false positive crosslinks, which is arguably the best result for this dataset as all other tools either substantially underestimate the FDR (MaxLynx, pLink, xiSearch, XlinkX) or detect a lot less crosslinks (MeroX). MaxLynx reports 10 more true positive hits than MS Annika on average, however at the cost of also 10 more false positive crosslinks. MeroX identifies only a single false positive crosslink on average but since it overall detects rather low numbers of crosslinks at a mean of 15 true positive hits, the average experimentally validated FDR is still above target at 4.23%. The search engine pLink yields a similar result to MaxLynx reporting 103 true positive and 14 false positive crosslinks on average across the three replicates with an experimentally validated FDR of 11.85%. Performing slightly better than pLink is xiSearch with an average of 120 unique true positive crosslink identifications and a mean experimentally validated FDR of 11.24%, resulting from 15 false positive hits on average. The search engine XlinkX reports the worst result also for this dataset, identifying 92 true positive and 36 false positive unique crosslinks on average, severely underestimating the FDR at an average experimentally validated FDR of 27.89% which effectively means that every fourth identified crosslink is a false positive hit. Moreover, XlinkX failed to search the first of the three replicates due to a recurring arithmetic overflow error, therefore the presented results are averages from replicate two and three. In Supplementary Figure 5 we again show the intersection and union of results from MS Annika, MaxLynx and pLink for all three replicates including calculated experimentally validated FDRs. Agreement for this dataset is not as high as for the dataset by Beveridge and co-workers, noticeably also in a higher error rate among intersections, yielding up to two false positive hits per replicate that are reported by all three search engines.

**Fig. 5.**
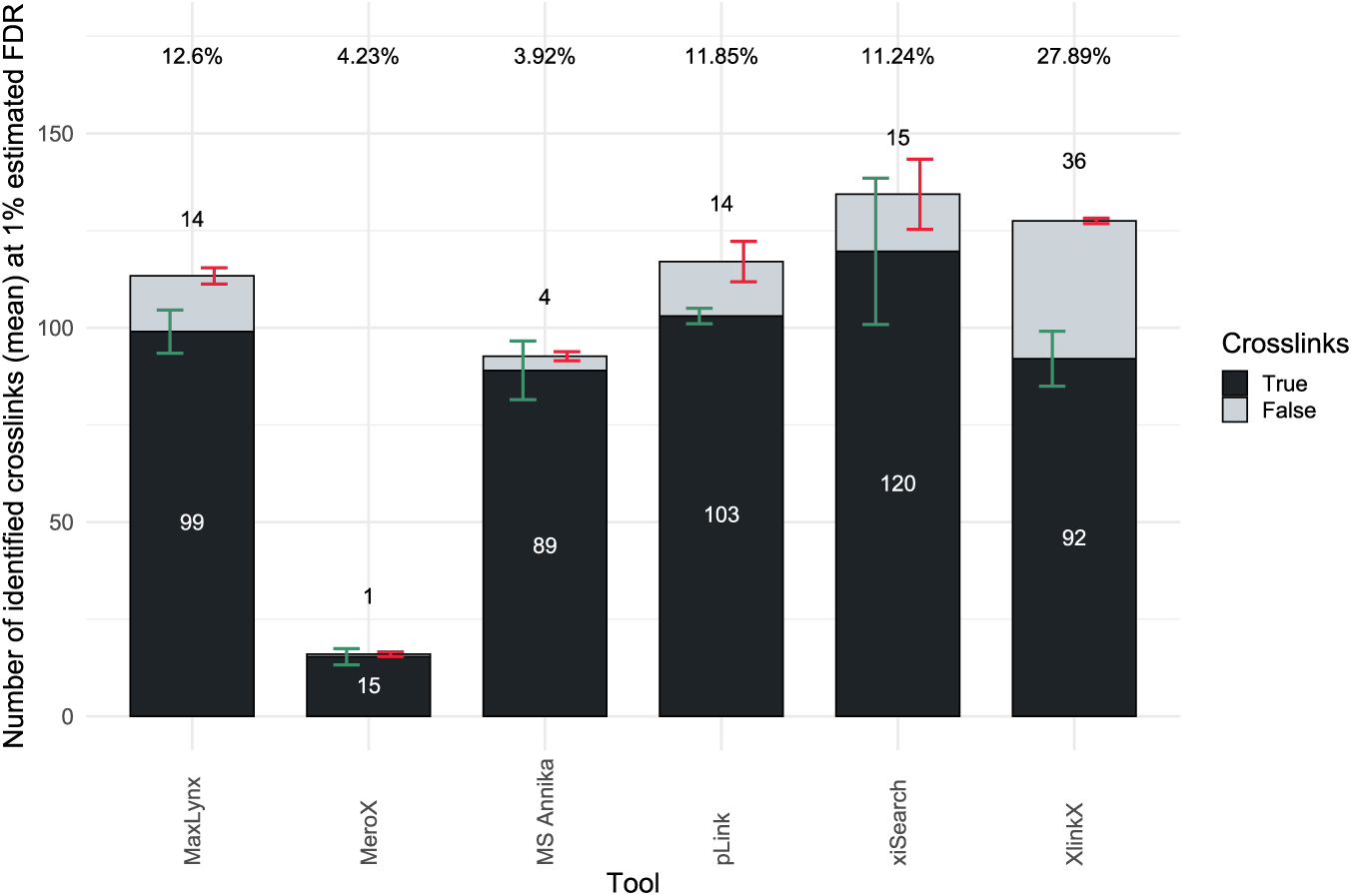
Identified crosslinks and experimentally validated FDRs of the different crosslinking search engines using the benchmark dataset by Matzinger and co-workers where synthetic peptides of their acidic library were crosslinked in seperate groups with the crosslinker ADH [46]. MS Annika reports the lowest experimentally validated FDR at 3.92%, identifying 89 true positive and 4 false positive crosslinks on average. MaxLynx [6], pLink [50] and xiSearch [51] detect more true positive crosslinks but also severely underestimate FDR, reporting experimentally validated FDRs of more than 10% above the 1% target. XlinkX [52] shows the highest experimentally validated FDR at 27.89%, while MeroX [48, 49] reports the lowest number of identified true positive crosslinks at an average of 15. Results were validated for 1% estimated FDR. All numbers are averages from three technical replicates (n = 3) except for XlinkX which failed to search replicate one (n = 2), the number of crosslinks was rounded to the closest integer value. Percentage numbers above the bars denote the average calculated experimentally validated FDR rounded to two decimal places. Error bars denote the standard deviation, in green for true positive crosslinks and in red for false positive crosslinks.

### 2.5 Diagnostic ions are not sufficient for crosslink spectrum detection

Steigenberger and co-workers suggest the usage of diagnostic ions for distinguishing mass spectra that contain crosslinked species from mass spectra that do not contain them [53]. Due to the complexity of non-cleavable crosslink searches it would be highly beneficial to be able to filter out mass spectra that do not contain crosslinked peptides which therefore would avoid spending computational resources on searching a spectrum with no valid result. We investigated the usage of diagnostic ions for our noncleavable search algorithm and again used the dataset by Beveridge and co-workers for reference [45] where peptides were crosslinked with DSS [21]. However, contrary to the results reported in the publication by Steigenberger and co-workers, we observed a severe drop in true positive crosslink identifications when only searching mass spectra that contained at least one diagnostic ion as shown in Supplementary Figure 6. On average across the three replicates, 118 less unique true positive crosslinks are identified at 1% estimated FDR when searching only mass spectra with at least one diagnostic ion compared to searching all mass spectra. This constitutes an overall worse result as the number of false positive identifications does not change, therefore yielding a higher experimentally validated FDR of 2.62%. Furthermore, only about 19.5% of mass spectra contain diagnostic ions (exact numbers given in Supplementary Table 3), substantially speeding up the search process, however at a cost in result quality which we do not deem worth it. Nevertheless, if the used non-cleavable crosslinker gives raise to diagnostic ions at a sufficient frequency that allows efficient distinction between crosslinked and non-crosslinked spectra, diagnostic ions can be specified in MS Annika to be considered for search, but by default MS Annika searches all MS2 spectra.

### 2.6 Validating MS Annika results with xiFDR boosts identifications and provides better FDR estimation

Validation of crosslinking results has been a widely discussed topic in the crosslinking community ever since its inception, with no clear consensus on how to perform proper validation. Most crosslinking search engines provide their own validation tools that range from simple peptide-spectrum-match (PSM) validation, as for example in pLink [50], to more refined approaches that can even validate on protein-protein interaction level, as in xiSearch [51] with xiFDR [24]. MS Annika follows a very strict validation approach where results can be either validated at crosslink-spectrum-match (CSM) or crosslink level, both using a target-decoy approach [5]. In Section 2.4 we show that this strategy works very well for estimating FDR, however, for larger studies and protein databases a more sophisticated approach might be beneficial to improve MS Annika results. In that regard we explored integrating the tool xiFDR [24] into our crosslink identification workflow which handles validation of MS Annika results. xiFDR allows a more nuanced control over validation and is able to boost the number of crosslink identifications by accounting for different crosslink or protein groups while keeping the overall FDR constant. We show the applicability of xiFDR with MS Annika using a dataset by Lenz and co-workers that allows calculation of an experimentally validated FDR for inter crosslinks [54]. The dataset consists of over 2.1 million mass spectra of proteins from *E. coli* which were crosslinked with BS3 [22]. Moreover, it is known which proteins are able to interact and therefore inter crosslinks can be assessed for their validity depending on if they form a possible protein-protein interaction or not. Mass spectra were searched with MS Annika against the full *E. coli* proteome (n = 4350) as provided by the authors of the dataset via the ProteomeXchange [55] partner repository JPOSTrepo [56] with accession codes PXD019120 and JPST000845. Supplementary Figure 7 depicts a comparison of results using either MS Annika with the built-in FDR validation or MS Annika with validation by xiFDR: using xiFDR for crosslink validation not only boosts the total number of identified crosslinks from 5134 to 6594 but also lowers the experimentally validated FDR for inter crosslinks from 3.1% to 0.42%, reporting only three crosslinks that constitute a protein-protein interaction that is not valid. This demonstrates the advantage of using more sophisticated validation approaches like xiFDR for larger studies and protein databases, enhancing results by reporting more crosslinks with less false positive hits.

## 3 Discussion

The algorithm presented here is an efficient and robust solution for identification of non-cleavable crosslinks in up to proteome-wide studies that runs smoothly on commodity hardware. Even though technically pLink [50] is also able to search large proteome-wide experiments, our results show that pLink suffers from underestimation of FDR, reporting substantially more false positive identifications than permissible even for less complex samples. Moreover, it should be noted that all the other search engines evaluated within this study are not capable of analysing crosslinking experiments that need to consider protein databases of more than a couple of thousand proteins. In contrast, our algorithm suffers none of these drawbacks and shows high numbers of crosslink identifications while keeping FDR and search times low. We postulate that this will not only enable researchers to perform large scale experiments with non-cleavable crosslinkers that were previously unfeasible, but also allows reanalysis of the vast amount of already published crosslinking data with bigger protein databases, potentially uncovering new protein interactions and biological insights.

Furthermore, the implemented approach of using sparse matrix multiplication for candidate selection is a transferable solution for large search space problems where theoretical ions need to be matched against experimental mass spectra which occur in other areas of proteomics [57], as well as metabolomics [58] and lipidomics [59]. The design of a scoring function purely based on sparse matrix operations proved to be highly efficient in both time and memory complexity which causes it to be a compelling method for large problems. Additionally, the memory requirements of sparse matrices do not grow with their dimensions but rather with the number of non-zero elements, in theory enabling scoring functions of almost infinite precision as binning windows can be made arbitrarily small. Another advantage are the on-going developments and optimizations of sparse matrix multiplication [60] potentially making this approach even more attractive in the future. Finally, we propose that new and more sophisticated scoring functions for database search could be built using sparse tensors such as implemented in TensorFlow [61, 62] which are similarly optimised but would allow incorporation of additional features like peak intensity for scoring, effectively improving score quality and better reflecting how good a match between a peptide and spectrum is.

Moreover, to show the applicability of our new non-cleavable search, we crosslinked *C. elegans* nuclei and performed a proteome-wide search on the measured mass spectrometry data. The identified crosslinks allowed us to conduct a comprehensive structural analysis of the Box C/D RNP complex by combining the crosslinking results with structural modelling: we could confirm interaction of nol-58 with nol-56 and fib-1 as well as interaction between nol-56 and M28.5 which facilitate the formation of a stable and functional RNP complex. The AlphaFold2 [38–41] predicted three-dimensional structure showed clear violation of PPI links exceeding the maximum allowed crosslink distance of DSG. We refined the structure with AlphaLink2 [43, 44] incorporating the identified crosslink restraints which resulted in a better structural model, both in terms of higher ipTM score as well as in reduction in the number of crosslink distance violations. Despite the refinement, two crosslinks still exceeded the distance limit due to the inherent symmetry of the complex and limitations in the number of complex members for structure prediction. Nonetheless, our structural analysis offers a detailed and improved view of the Box C/D RNP complex in *C. elegans* and significantly enhances our understanding of the assembly and functional dynamics of the complex.

## 4 Methods

### 4.1 Implementation of a novel non-cleavabe search algorithm in MS Annika

The general idea of the new non-cleavable search algorithm in MS Annika is a two step approach: first, identify one of the two peptides (from hereon denoted as alpha peptide), and second, identify the complementary peptide (from hereon denoted as beta peptide) that makes up the complete crosslink. The second step is a trivial problem as the mass of the beta peptide can be inferred from the precursor mass of the spectrum, the mass of the alpha peptide, and the mass of the crosslinker as shown in Equation 1.

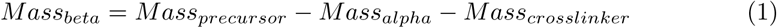

The identification of the alpha peptide is more challenging: as there is no information about the mass of the peptide available, all peptides in the database have to be considered as candidates for each spectrum. For large protein databases the number of candidate peptides easily reaches several millions, especially when decoys have to be considered. Therefore a search algorithm is needed that is able to efficiently score several million peptides in reasonable time. Over the last two decades computational vector and matrix operations have seen continuous improvement, ultimately giving rise to the now widely spread use of artificial neural networks and similar machine learning approaches in all areas of life which in turn further drove optimization [63]. The time efficiency of vector and matrix operations triggered us to design a search approach that is purely based on vector and matrix multiplications, however there was one problem that still remained: with potentially millions of peptide candidates the encoding matrix would grow to an enormous size that would be impossible to store in memory. Nonetheless, since most of the elements in the encoding matrix are zero, we explored the usage of sparse matrices which drastically reduced the memory footprint and allowed us to save complete encoded databases of even proteome-wide studies in memory. In the final implementation peptides are encoded as sparse vectors and mass spectra are either encoded as dense or sparse vectors, depending on the algorithm, which is a user-definable parameter.

The idea of encoding mass spectra as vectors or matrices has been previously explored for fast calculation of the cross-correlation score in Comet [64, 65] or for spectral library search using an approximate nearest neighbour approach in ANNSOLO [66]. Comet encodes mass spectra as sparse matrices by binning peaks in very small m/z windows where the matrix index corresponds to the m/z window and the matrix value to the observed intensity. Similarly, ANN-SOLO bins peaks into vectors also applying very small m/z windows where the vector index indicates the m/z window and the vector value the observed intensity. Moreover, ANN-SOLO also hashes the encoding vectors to reduce the vector dimensions and speed up the approximate nearest neighbour search. In MS Annika we employ an analogous approach, however with two significant changes: firstly, peaks are modelled as gaussian distributions with a mean corresponding to the peak’s m/z value and a standard deviation equal to *tolerance*/3, where tolerance is user-definable parameter and has to be given in Dalton. The gaussian peaks are then binned into vector indices using 0.01 m/z windows. Secondly, instead of using the peak intensity for the values of the vector, the values are given by the probability density function of the gaussian distribution of each peak as described in Equations 2 - 4. In Equation 2 the parameter y denotes the vector value and x the vector index while µ and σ are given in Equation 3 and 4, respectively.

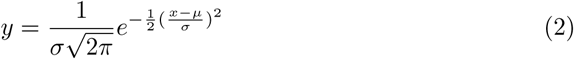

where

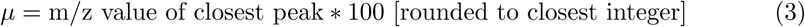

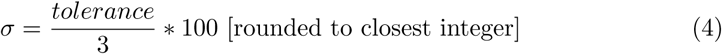

The idea is that experimentally measured peaks have to be considered with a certain tolerance to account for instrument errors and we postulate that errors are normally distributed with smaller errors being more likely than larger errors which is modelled by the gaussian distribution. Choosing a standard deviation of *tolerance*/3 denotes that more than 99% of the errors are within the instrument’s tolerance. Summarising, every mass spectrum can be encoded as a single sparse float vector and since we only consider peaks up to 5000 m/z the dimensionality of such a vector is 500 000. Supplementary Section 1 and Supplementary Figure 8 describe the spectrum encoding in more detail including pseudo-code and graphical explanation.

MS Annika goes one step further by also representing the protein database that is used for search as a sparse matrix: after *in silico* digestion of the proteins into peptides, all m/z values of the theoretical ions for each peptide are calculated and each peptide is encoded as a vector by binning the theoretical ion m/z values into vector indices using 0.01 m/z windows. The vector values for the given indices are either all one or one divided by the peptide’s length if the user wishes to normalise for peptide length. As a result each peptide in the database is represented as a sparse float vector with 500 000 dimensions, as again we do not consider theoretical ions beyond 5000 m/z. The whole database can therefore be encoded as an *M* x 500 000 sparse float matrix where M is the number of peptides in the database. More in-depth examples illustrating the peptide encoding are given in Supplementary Section 1 and Supplementary Figure 9. MS Annika uses this representation to calculate an approximate score for each peptide for a given mass spectrum to find likely candidates for crosslink identification. The approximate score for a peptide p given a mass spectrum s is calculated as shown in Equation 5: in brief the score is the dot product of the encoding vector of p and the encoding vector of s. This score can be interpreted as a measure of correlation between the ion series of p and the peaks of the experimental mass spectrum s. In the simplest case when the gaussian peak modelling for the spectrum encoding is disabled, the score represents exactly the number of matched peaks between p and s. Enabling gaussian peak modelling gives a deviation of this score, where higher scores denote that peaks were matched with higher precision. Optionally MS Annika normalises this score by peptide length, as described in the peptide vector encoding.

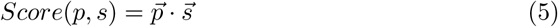

This approach can be easily extended to score all peptides against one or more given mass spectra by using all peptide encoding vectors (therefore a matrix) where the problem then can be denoted as simple matrix multiplication. In order to calculate the scores for all peptides in the database for all mass spectra, Equation 5 can be rewritten as in Equation 6. Essentially p and s become matrices instead of vectors, ^P⃗^ is the sparse encoding matrix of all peptides P in the database while ^S⃗^ is the encoding matrix of all mass spectra S. Scores(P, S) becomes an *M* x *N*-dimensional matrix of all scores Score(p, s) where *M* is the number of peptides in the database and *N* is the number of mass spectra.

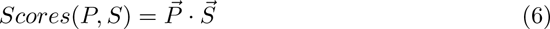

In total MS Annika implements 11 different algorithms to calculate Equation 6 out of which eight run on the CPU using the linear algebra library Eigen [67] which leverages modern CPU instruction sets for fast matrix operations and three run on the GPU using Nvidia CUDA cuSPARSE. Table 1 lists all algorithms available in MS Annika and Supplementary Figure 10 and 11 show an overview of the performance for each algorithm. The choice of algorithm is up to the user and depends on the available hardware, by default MS Annika uses algorithm *i32CPU DM* which works on all systems and which was used for all results presented in this study. All algorithms are implemented in the C++ programming language.

**Table 1.** Overview of the different matrix multiplication algorithms implemented and available in the MS Annika non-cleavable search. For algorithms using a vector-sbased spectrum encoding the multiplication is repeated *n* times, where *n* is the number of mass spectra. Usually it is impossible to score all mass spectra in a single computation as the resulting matrix would not fit in memory, therefore matrix-based spectrum encoding algorithms batch the computation using batches of 100 mass spectra per computation. For integer compute types the original float values are multiplied by 1000 and then rounded to the closest integer.

| Name | Compute Type | Peptide Encoding | Spectrum Encoding | Device |
| --- | --- | --- | --- | --- |
| f32CPU_SV | 32-bit Float | Sparse Matrix | Sparse Vector | CPU |
| i32CPU_SV | 32-bit Integer | Sparse Matrix | Sparse Vector | CPU |
| f32CPU_SM | 32-bit Float | Sparse Matrix | Sparse Matrix | CPU |
| i32CPU_SM | 32-bit Integer | Sparse Matrix | Sparse Matrix | CPU |
| f32CPU_DV | 32-bit Float | Sparse Matrix | Dense Vector | CPU |
| i32CPU_DV | 32-bit Integer | Sparse Matrix | Dense Vector | CPU |
| f32CPU_DM | 32-bit Float | Sparse Matrix | Dense Matrix | CPU |
| i32CPU_DM | 32-bit Integer | Sparse Matrix | Dense Matrix | CPU |
| f32GPU_DV | 32-bit Float | Sparse Matrix | Dense Vector | GPU |
| f32GPU_DM | 32-bit Float | Sparse Matrix | Dense Matrix | GPU |
| f32GPU_SM | 32-bit Float | Sparse Matrix | Sparse Matrix | GPU |

For optimal computational performance MS Annika by default scores all peptides against multiple mass spectra at once because sparse matrix * matrix multiplication is generally faster than multiple sparse matrix * vector multiplications (as presented in Supplementary Figure 11). Using sparse matrix multiplication for approximating scores is extremely efficient both in terms of computational speed as well as in memory consumption: calculating the approximate scores for all peptides of the human SwissProt proteome (including decoys, ~ 4.4 million peptides) for one spectrum takes less than a second on a quad core mobile CPU while storing the complete sparse matrix of peptide candidates requires less than 8 GB of memory, allowing for proteome-wide searches on normal office laptops and other standard commodity hardware.

Subsequently, when the computation of these approximate scores for all peptides for a given mass spectrum is finished, the scores are used to take the top *N* peptide candidates to consider for crosslink identification, where *N* is user-definable parameter which by default is 100. MS Annika generates every possible peptidoform for each of the top peptide candidates and our in-house developed peptide search engine MS Amanda [36, 37] is used to calculate a more sophisticated score for each peptidoform. Even though multi-step search approaches are quite common in the field of computational proteomics and especially in crosslinking search engines [68], it has been argued that multi-step approaches might hinder robust FDR estimation as decoy peptides might not pass at the same rate as target peptides [69]. In MS Annika this problem is avoided as candidate peptides are not directly provided to MS Amanda but rather only their mass is given for identification. This ensures that even at the second step where peptidoforms are accurately scored, the number of potential decoy candidates is the same as the number of potential target candidates. The result of scoring with MS Amanda is a list of peptidoform candidates with a score highly reflective of match quality. MS Annika takes the top *M* peptidoforms of this list, where *M* again is a user-definable parameter and by default 10, and considers them to be possible alpha peptides for crosslink search. Furthermore, MS Annika then tries to identify complementary beta peptides that would make up a complete crosslink. As noted above in Equation 1, this is a trivial problem as the mass of the beta peptide can be easily calculated by subtracting the mass of the alpha peptide and the mass of the crosslinker from the precursor of the mass spectrum.

Finally, if any beta peptides are identified, crosslink-spectrum-matches (CSMs) are constructed for any combination of alpha and beta peptides that match the total precursor mass of the mass spectrum. The score of the CSM is the minimum score of the two peptides - this is identical to the cleavable search that we published previously [5]. The remaining steps of the search are all equal to the cleavable search: the CSM with the highest score is reported and used for validation, multiple CSMs denoting the same crosslink are grouped and the score of the crosslink is the maximum score of all CSMs. CSMs and crosslinks are validated using a target-decoy approach where target-target hits are considered as targets and target-decoy, decoy-target and decoy-decoy hits are considered as decoys.

### 4.2 Nuclei isolation from *C. elegans*

Worms were maintained at 20°C on Nematode Growth Medium (NGM) agar plates seeded with *Escherichia coli* OP50 [70]. Hermaphrodites were used in all experiments unless otherwise stated. The strain *M01E11.3(jf92[M01E11.3::unc-119(+)])I; jfsi38[gfp::rmh-1;cb-unc-119(+)]II; unc-119(ed3)III;vieSi146(pAD860;pCFJ151 ppie1::SV40::vhh4GFP4::TurboID::tbb-2UTR; cb unc-119(+))IV* was used for nuclei isolation and crosslinking experiments.

Worm nuclei extraction was performed as previously described [70, 71]. For this, one-sixth of a starved 60 mm plate was transferred onto a 100 mm plate seeded with freshly grown OP50 *E. coli* and worms were grown at 20°C for 3 days. Worms were then collected in M9 buffer and washed at least three times (sedimented by gravity at room temperature) to remove most of the OP50 bacteria. The final worm pellet was frozen at −80°C in 3 ml NP buffer (10 mM HEPES–KOH pH 7.6, 1 mM EGTA, 10 mM KCl, 1.5 mM MgCl_2_, 0.25 mM sucrose, 1 mM PMSF) containing Protease Inhibitor Cocktail (Roche). A 1 ml worm pellet obtained from 30–40 OP50-seeded 100 mm NGM plates was used for fractionation. To isolate the nuclei, worms were disrupted using a cooled metal Wheaton tissue grinder and the suspension was filtered using first a 100 µm mesh and then a 40 µm mesh. The filtered suspension was clarified at 300 g for 2 min at 4°C, and the supernatant from this step, containing the nuclei, was centrifuged at 2500 g for 10 min at 4°C. Now this supernatant contained the cytosolic fraction and the germline nuclei in the pellet.

### 4.3 Crosslinking procedure

Germline nuclei were resuspended in crosslinking buffer (50mM HEPES-KOH pH 7.6) and crosslinked using disuccinimidyl glutarate (DSG, Cat-no.: 20593 Thermo Scientific) at a concentration of 3 mM for 45 min on ice. The crosslink reaction was quenched by adding 100 mM Tris (pH 7.4) and incubating it for 5 minutes at room temperature.

### 4.4 In-solution digest

The isolated and crosslinked nuclei were incubated in guanidine hydrochloride with a final concentration of 8 M and 1% ProteaseMax (Cat-no.: V2071, Promega). The solution was sonicated for 30 seconds at an amplitude of 80% with a 0.5-second cycle, followed by chilling on ice; this process was repeated three times. Proteins were reduced using 10 mM DTT, following a 30-minute incubation period at 50°C and alkylated using 50 mM IAA for 30 minutes at room temperature in the dark. The sample was diluted to 2 M guanidine hydrochloride using 50 mM HEPES, pH 7.3. Subsequently, 1 mM MgCl2 and 25 U/L Benzonase were added to digest DNA and RNA, and the mixture was incubated for 1 hour at 37°C. Afterwards, LysC was added to a final concentration of 3 ng/µL, and the mixture was incubated for another hour at 37°C. Trypsin was added at a ratio of 1:100 (enzyme to protein) for final digestion and the sample was incubated overnight at 37°C. Finally, the sample was acidified using 10% TFA to reach a final concentration of 0.5% and hence remove the ProteaseMax from the sample by precipitation.

Digested peptides were desalted using C18 columns (Sep-Pak C18 1 cc Vac Cartridges, waters). The column material was activated by flushing the column once with methanol, following equilibration using 0.1% trifluoroacetic acid (TFA) until all traces of MeOH were washed away. The sample, adjusted to a pH of 3, was then loaded onto the column. Following sample loading, the column was washed three times with 0.1% TFA and subsequently eluted using 80% acetonitrile (ACN) in 0.1% TFA. The ACN content in the eluate was removed using a SpeedVac, and the sample was lyophilized.

### 4.5 Size exclusion fractionation

Purified samples were reconstituted in 0.1% TFA to a final concentration of 3 µg/µL. 60µg of peptides were injected per sample and condition on a Dionex UltiMate 3000 HPLC system (Thermo Fisher Scientific) consisting of autosampler, SD-pumps, and UV detectors. Fractions had to be collected manually. Peptides were separated on a TSKgel SuperSW2000 column (4.6 mm ID x 30 cm L, P/N: 0018674, Tosoh Bioscience) at a flow rate of 300 µL min-1 using the SEC mobile phase (30% ACN in 0.1% TFA) and an isocratic gradient. The separation was monitored by UV absorption at 214 nm. half-a-minute fractions (150 µl) were collected into 0.6 µL low-bind reaction tubes over a separation window of 6 min. For analysis by liquid chromatography (LC)-MS/MS, fractions of interest (retention times 6-12 min) were removed and evaporated to dryness.

### 4.6 Mass spectrometry analysis

LC-MS/MS analysis was performed using an Orbitrap Exploris 480 with Field asymmetric ion mobility spectrometry (FAIMS) interface (Thermo Fisher Scientific, Waltham, Massachusetts, United States) coupled with a Vanquish Neo HPLC system (Thermo Fisher Scientific, Waltham, Massachusetts, United States). A trap column PepMap C18 (5 mm × 300 µm ID, 5 µm particles, 100 Å pore size) (Thermo Fisher Scientific, Waltham, Massachusetts, United States) and an analytical column PepMap C18 (500 mm × 75 µm ID, 2 µm, 100 Å) (Thermo Fisher Scientific, Waltham, Massachusetts, United States) were employed for separation. The column temperature was set to 50°C. Sample loading was performed using 0.1% trifluoroacetic acid in water with a flow rate of 25 µL/min for 3 min. Mobile phases used for separation were as follows: (A) 0.1% formic acid (FA) in water; (B) 80% acetonitrile, 0.1% FA in water. Peptides were eluted using a flow rate of 230 nL/min, with the following gradient: from 2% to 37% phase B in 80 min, 37% to 47% phase B in 7 min, from 47% to 95% phase B in 3 min, followed by a washing step at 95% for 5 min, and re-equilibration of the column. FAIMS separation was performed with the following settings: inner and outer electrode temperatures were 100°C, FAIMS carrier gas flow was 4.2 L/min, compensation voltages (CVs) of −50, −60, and −70 V were used in a stepwise mode during the analysis. The mass spectrometer was operated in a data-dependent mode with cycle time 2s, using the following full scan parameters: m/z range 350-1600, nominal resolution of 120 000, with an automated gain control (AGC) target set to standard, and 90 ms maximum injection time. For higher-energy collision-induced dissociation (HCD) MS/MS scans, a stepped normalised collision energy (NCE) of 25%; 27%; 33% and MS2 resolution of 30 000 was used. Precursor ions were isolated in a 2 Th window with no offset and accumulated for a maximum of 70 ms or until the AGC target of 200% was reached. Precursors of charge states from 2+ to 6+ were scheduled for fragmentation. Previously targeted precursors were dynamically excluded from fragmentation for 15 seconds. The sample load was 500 ng. Detailed parameters can be found in each raw file under the instrument method section.

### 4.7 Construction of the filtered *C. elegans* protein database

In order to study the influence of protein database size on the non-cleavable search in MS Annika we used two different databases for crosslink identification: 1) the full *C. elegans* proteome and 2) a filtered proteome only containing the most abundant proteins found in a preliminary linear search. The filtered proteome was constructed as follows: mass spectrometry RAW files were loaded in Proteome Discoverer 3.1 (version 3.1.0.638) and mass spectra were deisotoped and charge deconvoluted with the IMP MS2 Spectrum Processor node. Mass spectra were then searched with MS Amanda 3.0 [36, 37] (version 3.1.21.532) using the full *C. elegans* reference proteome (n = 26 695, UniProt Proteome ID UP000001940, retrieved 15. March 2024). The digestion enzyme was set to trypsin with a maximum of 3 missed cleavages allowed. The minimum peptide length was set to 5 and the maximum peptide length to 30 amino acids. Precursor mass tolerance was set to 5 ppm and fragment mass tolerance to 10 ppm. Carbamidomethylation of cysteine was considered as a fixed modification and oxidation of methionine, phosphorylation of serine, threonine and tyrosine, deamidation of asparagine and glutamine, carbamylation of methionine, acetylation of the protein nterminus, as well as modification of lysine by the monolink forms of DSG were specified as possible variable modifications. Results were validated with Percolator [72] (version 3.05.0). The filtered protein database was then constructed by selecting proteins that had more than two high-confidence (1% FDR) PSMs associated in at least one of the three biological replicates. The final filtered protein database was exported to fasta format and consisted of 3069 proteins. Construction of the database was done with an in-house developed python script using biopython [73].

### 4.8 Crosslink identification and validation of *C. elegans*

Mass spectrometry RAW files were loaded in Proteome Discoverer 3.1 (version 3.1.0.638) and searched with our standard crosslink identification workflow for large studies: mass spectra were deisotoped with the IMP MS2 Spectrum Processor node and then searched for linear and monolinked peptides with MS Amanda 3.0 [36, 37] (version 3.1.21.532) using either the full *C. elegans* reference proteome (n = 26 695, UniProt Proteome ID UP000001940, retrieved 15. March 2024) or a filtered version as described in Section 4.7. Trypsin was specified as the digesting enzyme and the maximum number of allowed missed cleavages was set to 3. The minimum peptide length was again set to 5 and the maximum peptide length to 30 amino acids. For identification the precursor mass tolerance was set to 5 ppm and the fragment mass tolerance for matching was set to 10 ppm. Carbamidomethylation of cysteine was defined as a fixed modification and oxidation of methionine, phosphorylation of serine, threonine and tyrosine, deamidation of asparagine and glutamine, carbamylation of methionine, acetylation of the protein n-terminus, as well as modification of lysine by the monolink forms of DSG were considered as variable modifications. After linear and monolinked peptide identification, results were validated with a standard target-decoy approach and any mass spectrum with a high-confidence PSM (1% FDR) was filtered out and not considered for crosslink search. Crosslinks were identified with MS Annika 3.0 (version 3.0.1) using the non-cleavable search approach with the same protein database as in MS Amanda. The digestion enzyme was again set to trypsin with a maximum of 3 missed cleavages. For crosslinked peptides the minimum considered peptide length was again 5 amino acids and the maximum 30 amino acids. Again a precursor mass tolerance of 5 ppm and fragment mass tolerance of 10 ppm were used. The crosslinker parameter was set to DSG with allowed reactions to lysine and the protein n-terminus. Carbamidomethylation of cysteine was set as a fixed modification and oxidation of methionine as a variable modification. The top 2 alpha peptides were considered for CSM creation. Finally, after search, MS Annika CSMs were exported to xiFDR [24] format using a newly developed MS Annika to xiFDR exporter script and validated with xiFDR (version 2.2.1) using 1% crosslink FDR with boosting enabled. Supplementary Table 4 gives a summary of all search and validation parameters.

### 4.9 AlphaFold2 Multimer screening

AlphaFold2 Multimer [38–41] was used to predict interactions between nol-58 and 29 proteins putative nol-58 interactors based on known interactions extracted from the literature. We employed a custom script to run pairwise predictions on a local CPU and GPU cluster, using MMseqs (git@92deb92) for local Multiple Sequence Alignment (MSA) creation and colabfold (git@7227d4c) for structure prediction with 5 models per prediction and omitting structure relaxation. Predictions with an average iPTM score of >0.6 were considered putative hits and diagnostic plots (PAE plot, pLDDT plot and sequence coverage) as well as the generated structures were manually inspected.

After selecting the top 5 hits, the prediction of the complex was performed by running pairwise comparisons of nol-58, NEDG-01330, fib-1 comprised in one fasta as chain A, chain B and chain C respectively against M28.5 and nol-56 in a second fasta. The crosslink restrains from the crosslinking experiment were plotted onto the Rank 1 structure (ipTM 0.725). Diagnostic plots for this complex prediction like PAE plot, pLDDT plot and sequence coverage are shown in Supplementary Figure 12.

### 4.10 AlphaLink2 structure refinement

The AlphaLink2 structure refinement was performed as described in Stahl et al. [44] with default settings and following the instructions for container implementation on a cluster system on GitHub (https://github.com/lhatsk/AlphaLink and https://github.com/Rappsilber-Laboratory/AlphaLink2). In short, extracted from Stahl *et al.*, OpenFold was enhanced by adding a crosslink embedding layer to map contact maps or distograms into the 128-dimensional z-space of AlphaFold2/OpenFold, integrated into the pair representation (z), along with a group embedding layer for ambiguous crosslinks. MSAs were randomly subsampled each epoch to achieve N*_eff_* between 1 and 25, reflecting non-redundant sequences below 80% identity. Using AlphaFold2 2.0 weights, the network was refined on 13 000 proteins from the trRosetta training set with simulated crosslinking data, using OpenFold v0.1.0 and model 5 ptm. UniRef90 v2020 01, MGnify v2018 12, Uniclust30 v2018 08, BFD, PDB (May 2020), and PDB70 (May 2020) were used to mimic CASP14 settings. For CAMEO, 45 targets released after AlphaFold2 were considered, excluding those with TM scores above 0.8. Network weights were downloaded from Zenodo (https://zenodo.org/records/8007238). The “AlphaLink-Multimer SDA v3.pt” parameter file was used for all predictions.

### 4.11 Workflow for the analysis of the benchmark dataset by Beveridge and co-workers

Data was retrieved from ProteomeXchange [55] via the PRIDE partner repository [74] with identifier PXD014337. Detailed information about the data can be found in the respective publication by Beveridge and co-workers [45]. In short, Beveridge and co-workers created a synthetic peptide library consisting of 95 peptides from *Streptococcus pyogenes* Cas9 that were divided into 12 groups and crosslinked within their groups using the crosslinker DSS [21]. Importantly, the premise is that any identified crosslink that is composed of two peptides of different groups or non-synthesized peptides is a false positive, allowing the computation of an experimentally validated FDR and comparison to the FDR estimation of the identifying crosslinking search engine. Samples were measured in technical triplicates on a Q Exactive HF-X (Thermo Fisher Scientific). For MeroX [48, 49] and xiSearch [51] RAW files were exported to MGF format using Proteome Discoverer 3.1 (version 3.1.0.638) since they do not support RAW file input, for all other search engines the RAW files were used directly. Mass spectrometry data was searched with MaxLynx [6] (part of MaxQuant [47], version 2.6.2.0), MeroX (version 2.0.1.4), MS Annika (version 3.0.1) in Proteome Discover 3.1 (version 3.1.0.638), pLink (version 2.3.11) [50], xiSearch (version 1.7.6.7) using xiFDR (version 2.2.1) [24] for validation, and XlinkX (version as distributed with the Proteome Discoverer third-party nodes installer) [52] in Proteome Discoverer 3.1 (version 3.1.0.638). For all search engines we used settings as given in the publication by Beveridge and co-workers: the considered protein database consisted of the sequence of *S. pyogenes* Cas9 and 10 contaminant proteins (n = 11), the digestion enzyme was set to trypsin with a maximum of 3 missed cleavages allowed and peptides with a length of at least 5 but not more than 60 amino acids were permissible for search. The precursor mass tolerance was set to 5 ppm and the fragment mass tolerance to 20 ppm. Carbamidomethylation of cysteine was applied as a fixed modification and oxidation of methionine was specified as a possible variable modification. The crosslinker parameter was set to DSS with reactions to lysine and the protein n-terminus allowed. Results were validated for 1% estimated FDR and subsequently analysed with IMP-X-FDR (version 1.1.0) [46] to calculate experimentally validated FDRs. For MaxLynx the crosslink search was set up in MaxQuant as described in their publication [6], for MS Annika the standard non-cleavable workflow was employed, and for XlinkX the non-cleavable HCD/CID MS2 Proteome Discoverer workflow was used for crosslink search. Supplementary Table 5 gives a summary of all search and validation parameters. Workflows for MS Annika and XlinkX are shown in Supplementary Figure 13 and 14, respectively.

### 4.12 Workflow for the analysis of the benchmark dataset by Matzinger and co-workers

Data was retrieved from ProteomeXchange [55] via the PRIDE partner repository [74] with identifier PXD029252. Detailed descriptions of the data are given in the respective publication by Matzinger and co-workers [46]. Summarising, Matzinger and co-workers engineered a synthetic peptide library that features a total of 141 peptides from 38 different proteins of the *Escherichia coli* ribosomal complex. Analogous to the synthetic peptide library by Beveridge and co-workers [45] the peptides were split into groups of 6 - 10 peptides each and crosslinked groupwise, facilitating the calculation of an experimentally validated FDR after identification since it is known which peptides can be crosslinked. Specifically, the assumption is again that any crosslink between peptides of different groups or non-synthesized peptides is a false positive. The analysed samples were crosslinked with ADH and measured in three technical replicates on a Q Exactive HF-X (Thermo Fisher Scientific). Mass spectrometry data was processed using the same tools and in the same way as described in 4.11, with the following deviations: the protein database used for search was composed of 171 sequences from the *E. coli* ribosomal complex, again using trypsin as the digestion enzyme with a maximum of 3 missed cleavages. The minimum peptide length was set to 6 and the maximum peptide length to 60 amino acids. Precursor mass tolerance and fragment mass tolerance were set to 5 ppm and 10 ppm respectively. Carbamidomethylation of cysteine was considered as a fixed modification and oxidation of methionine as a variable modification. The crosslinker parameter was specified as ADH with allowed reactions to aspartic acid and glutamic acid. Validation was set to 1% estimated FDR and the validated results were post-processed with the tool IMP-X-FDR (version 1.1.0) [46] to calculate experimentally validated FDRs. Search engine specific parameters were again set up according to developer recommendations, if available: MaxLynx [6] and MaxQuant [47] were configured as described in the MaxLynx publication [6], in MS Annika the standard non-cleavable workflow was employed with the addition of the IMP MS2 Spectrum Processor node as the sample was more complex, and for XlinkX the Proteome Discoverer workflow for non-cleavable HCD/CID MS2 was applied. Supplementary Table 6 presents a summary of all search and validation parameters. Furthermore, Supplementary Figure 14 and 15 explain the used Proteome Discoverer workflows for MS Annika and XlinkX.

### 4.13 Implementation of a xiFDR exporter and analysis of the dataset by Lenz and co-workers

MS Annika result tables were updated to include all necessary information required for validation with xiFDR [24]. Moreover, an exporter script was written in python that allows export of MS Annika CSMs to the specific CSV format required by xiFDR which is described in the xiFDR documentation on GitHub (https://github.com/Rappsilber-Laboratory/xiFDR). In order to validate the applicability of MS Annika with xiFDR, data from a study by Lenz and co-workers [54] that allows calculation of an experimentally validated inter crosslink FDR was retrieved from ProteomeXchange [55] via the JPOSTrepo partner repository [56] with accession codes PXD019120 and JPST000845. An in-depth explanation of the study is given in the respective publication [54]. In brief, Lenz and co-workers separated *E. coli* cell lysate using size exclusion chromatography which resulted in 44 fractions with molecular weight ranging from ~3 MDa to 150 kDa. Part of each fraction was used to create elution profiles of each protein across all 44 fractions using label-free quantitative proteomics. The remainder of each fraction was crosslinked with BS3 [22] and finally all fractions were pooled and analysed using LC-MS. The presumption for calculating experimentally validated inter crosslink FDRs is that only proteins which eluted in the same size exclusion fraction may be crosslinked, and contrary any crosslink consisting of peptides from two proteins that are not from the same fraction is a false positive. For our analysis the retrieved MGF files were loaded in Proteome Discoverer 3.1 (version 3.1.0.638) and mass spectrometry data (around 2.1 million mass spectra) was first searched with MS Amanda (version 3.1.21.532) [36, 37] to identify linear and monolinked peptides. Wherever possible we applied the settings recommended by Lenz and co-workers: the full *E. coli* proteome (n = 4350, retrieved from the dataset repository) was used for search, the digestion enzyme was set to trypsin with a maximum of 2 missed cleavages allowed. The minimum peptide length was specified as 6 and the maximum allowed peptide length was 60 amino acids. Precursor mass tolerance and fragment mass tolerance were set to 3 ppm and 5 ppm respectively. Carbamidomethylation of cysteine was considered as a fixed modification and oxidation of methionine as well as reaction of lysine with the monolink forms of BS3 were defined as variable modifications. Results of the MS Amanda search were validated for 1% estimated FDR using a standard target-decoy approach and mass spectra with an associated high-confidence (1% FDR) PSM were filtered out and not considered for crosslink search. The remaining mass spectra were searched with MS Annika using the same settings, except the mono-link forms of BS3 were not considered as variable modifications. Additionally, up to two missing precursor isotope peaks were allowed for identification and the crosslinker parameter was set as BS3 with possible reactions to lysine or the protein n-terminus. The top 2 alpha peptides were considered for CSM creation. Results were either validated with the built-in validation algorithm in MS Annika, or with xiFDR (version 2.2.1) using the exporter script described above. In both cases crosslinks were validated for 1% estimated FDR and boosting was enabled in xiFDR. The plausibility of inter crosslinks was assessed with an in-house developed script applying the rules outlined by Lenz and co-workers. Supplementary Table 7 summarises all parameters used during search and validation.

## 5 Data availability

All mass spectrometry proteomics data along with result files have been deposited to the ProteomeXchange consortium (http://proteomecentral.proteomexchange.org) [55] via the PRIDE partner repository [74] with the dataset identifier PXD055488. Result files of the re-analysis of the datasets by Beveridge and co-workers, Matzinger and coworkers and Lenz and co-workers have also been deposited to the ProteomeXchange consortium via the PRIDE partner repository with the dataset identifier PXD055512. All experimental metadata has been generated using lesSDRF [75].

## 6 Code availability

The newest version of MS Annika 3.0 including the new non-cleavable search is available free of charge for Proteome Discoverer 3.1 at https://ms.imp.ac.at/index.php?action=ms-annika or via the GitHub repository https://github.com/hgb-bin-proteomics/MSAnnika. The MS Annika 3.0 plugin node can be run with a free version of Proteome Discoverer, which is available for download from the Thermo Fisher website https://www.thermofisher.com/at/en/home/industrial/mass-spectrometry/liquid-chromatography-mass-spectrometrylc-ms/lc-ms-software/multi-omics-data-analysis/proteomediscoverer-software.html. MS Annika 3.0 makes use of the graphical user interface for workflow generation in Proteome Discoverer so users can easily set up searches with minimal bioinformatics knowledge. A detailed user manual, including descriptions of all tunable parameters, sample workflows and step-by-step instructions for running MS Annika 3.0, as well as licence information is also given in the MS Annika 3.0 repository. The candidate search algorithm, including encoding of peptides and mass spectra as sparse vectors, is - the same as MS Annika - written in C#, fully open source with a permissive MIT licence and available for all major platforms and architectures at https://github.com/hgb-bin-proteomics/CandidateSearch. The matrix multiplication backend is written in C++, also open source (MIT licence) and available for all major platforms and architectures via the repository https://github.com/hgb-bin-proteomics/CandidateVectorSearch. We also provide a template for custom matrix multiplication backends for researchers who want to take advantage of specific hardware, the template is available via https://github.com/hgb-bin-proteomics/CandidateVectorSearchtemplate. The python script for integrating xiFDR into MS Annika is available from the MS Annika exporters repository at https://github.com/hgb-bin-proteomics/MSAnnikaexporters. All scripts that were developed and used for data analysis and visualisation were deposited in https://github.com/hgb-bin-proteomics/MSAnnikaNCResults.

## 7 Author contributions

M.J.B. conceptualised, implemented and evaluated the non-cleavable search in MS Annika, performed data analysis and visualisation and wrote the manuscript. F.M. performed all *C. elegans* crosslinking experiments, performed data analysis and visualisation and wrote the manuscript. S.S.G. performed nuclei isolation of *C. elegans*. V.D. conceptualised and supervised the implementation and evaluation of the noncleavable search. M.M., K.M. and V.D. designed and supervised the study and revised the manuscript. All authors have given approval to the final version of the manuscript.

## 10 Ethics approval and consent to participate

Not applicable.

## 11 Consent for publication

Not applicable.

## 12 Materials availability

Not applicable.

## Supplementary information

The following supplementary information is provided:

- Supplementary Information.pdf:

1. Supplementary Section 1: Description of the encoding of mass spectra and peptides.
2. Supplementary Figure 1: Overlaps of identified crosslinks at 1% FDR in the *C. elegans* dataset using MS Annika for search and xiFDR [24] for validation.
3. Supplementary Figure 2: Influence of protein database size on the non-cleavable search of MS Annika when using the built-in valdiation algorithm.
4. Supplementary Figure 3: Overlaps of identified crosslinks at 1% FDR in the *C. elegans* dataset using MS Annika for search and validation.
5. Supplementary Table 1: Search times for crosslink identification in the three replicates of our *C. elegans* dataset using the full proteome-wide search.
6. Supplementary Table 2: Hardware configuration of the system that we used for benchmarking and for the more demanding proteome-wide searches.
7. Supplementary Table 3: Number of mass spectra containing diagnostic ions in the dataset by Beveridge and co-workers [45].
8. Supplementary Figure 4: Overlaps of identified crosslinks at 1% FDR in the dataset by Beveridge and co-workers [45].
9. Supplementary Figure 5: Overlaps of identified crosslinks at 1% FDR in the dataset by Matzinger and co-workers [46].
10. Supplementary Figure 6: Comparison of running the MS Annika non-cleavable search on either all spectra or only spectra containing diagnostic ions of the benchmark dataset by Beveridge and co-workers [45].
11. Supplementary Figure 7: Comparison of crosslink results of MS Annika using either the in-built FDR validation or xiFDR [24] for validation, as demonstrated on the dataset by Lenz and co-workers [54].
12. Supplementary Figure 8: Visualization of the mass spectrum encoding.
13. Supplementary Figure 9: Visualization of the peptide encoding.
14. Supplementary Figure 10: Synthetic benchmark of the different sparse matrix multiplication algorithms implemented in MS Annika.
15. Supplementary Figure 11: Benchmark of the different sparse matrix multiplication algorithms implemented in MS Annika using real data.
16. Supplementary Figure 12: Diagnostic plots for the Box C/D RNP complex prediction.
17. Supplementary Figure 13: The standard Proteome Discoverer workflow for crosslink identification with MS Annika.
18. Supplementary Figure 14: The non-cleavable HCD/CID MS2 Proteome Discoverer workflow for crosslink identification with XlinkX [52].
19. Supplementary Figure 15: The Proteome Discoverer workflow for crosslink identification with MS Annika for complex samples.
- Supplementary Table4.xlsx: Parameters used for searching the *C. elegans* data.
- Supplementary Table5.xlsx: Parameters used for searching the dataset by Beveridge and co-workers [45].
- Supplementary Table6.xlsx: Parameters used for searching the dataset by Matzinger and co-workers [46].
- Supplementary Table7.xlsx: Parameters used for searching the dataset by Lenz and co-workers [54].

## Acknowledgements

This work was supported by the F&E Infrastruk-turförderung 4. Ausschreibung 2022/01 (AT-SCP, https://projekte.ffg.at/projekt/4795911, accessed on 5 December 2023) of the Austrian Research Promotion Agency (FFG). This work was further funded by the project LS20-079 of the Vienna Science and Technology Fund and the project P35045-B of the Austrian Science Fund (FWF, Grant DOI 10.55776/P35045) and the the ESPRIT program project number ESP 566 (Grant-DOI 10.55776/ESP566). Work at the Max Perutz Labs (MPL) was funded by project SFB F 8805-B of the Austrian Science Fund (FWF). We thank the developers and community of the Eigen linear algebra library for their input on matrix multiplication and open sourcing their library. We further thank the developers of Nvidia CUDA cuSPARSE for providing this library. Our gratitude further goes to L. Buur, S. Dorl, J. Vetter and S. Winkler for fruitful discussions and their feedback on the manuscript. For the purpose of open access, the author has applied a CC BY public copyright licence to any Author Accepted Manuscript version arising from this submission.

