## Supplementary Information for "Proteome-wide non-cleavable crosslink identification with MS Annika 3.0 reveals the structure of the *C. elegans* Box C/D complex"

**Table of Contents**


- **Supplementary Section 1:** Encoding of mass spectra and peptides.
- **Supplementary Figure 1:** Overlaps of identified crosslinks at 1% FDR in the *C. elegans* dataset using MS Annika for search and xiFDR [1] for validation.
- **Supplementary Figure 2:** Influence of protein database size on the non-cleavable search of MS Annika when using the built-in valdiation algorithm.
- **Supplementary Figure 3:** Overlaps of identified crosslinks at 1% FDR in the *C. elegans* dataset using MS Annika for search and validation.
- **Supplementary Table 1:** Search times for crosslink identification in the three replicates of our *C. elegans* dataset using the full proteome-wide search.
- **Supplementary Table 2:** Hardware configuration of the system that we used for benchmarking and for the more demanding proteome-wide searches.
- **Supplementary Table 3:** Number of mass spectra containing diagnostic ions in the dataset by Beveridge and co-workers [2].
- **Supplementary Figure 4:** Overlaps of identified crosslinks at 1% FDR in the dataset by Beveridge and co-workers [2].
- **Supplementary Figure 5:** Overlaps of identified crosslinks at 1% FDR in the dataset by Matzinger and co-workers [3].
- **Supplementary Figure 6:** Comparison of running the MS Annika non-cleavable search on either all spectra or only spectra containing diagnostic ions of the benchmark dataset by Beveridge and co-workers [2].
- **Supplementary Figure 7:** Comparison of crosslink results of MS Annika using either the built-in FDR validation or xiFDR [1].
- **Supplementary Figure 8:** Visualization of the mass spectrum encoding.
- **Supplementary Figure 9:** Visualization of the peptide encoding.
- **Supplementary Figure 10:** Synthetic benchmark of the different sparse matrix multiplication algorithms implemented in MS Annika.
- **Supplementary Figure 11:** Benchmark of the different sparse matrix multiplication algorithms implemented in MS Annika using real data.
- **Supplementary Figure 12:** Diagnostic plots for the Box C/D RNP complex prediction.
- **Supplementary Figure 13:** The standard Proteome Discoverer workflow for crosslink identification with MS Annika.
- **Supplementary Figure 14:** The non-cleavable HCD/CID MS2 Proteome Discoverer workflow for crosslink identification with XlinkX [4].
- **Supplementary Figure 15:** The Proteome Discoverer workflow for crosslink identification with MS Annika for complex samples.

### 1 Encoding of mass spectra and peptides

In order to demonstrate the process of encoding mass spectra and peptides as vectors, consider the following:

1. Peptide $P$ which is a float array of calculated m/z values for all theoretical ions.
2. Integer number $p$ of theoretical ions in $P$.
3. Mass spectrum $S$ which is a float array of m/z values of its peaks. (The intensities are not used for this algorithm).
4. Integer number $s$ of peaks in $S$.
5. Tolerance $T$ which is single float value and is given in Dalton.

The mass spectrum encoding vector is calculated as follows (here implemented in the C++ language):

```
const int dims = 500000;
const double n = 0.3989422804014326779399460599343 8;
float normpdf(float x, float mu, float sigma) {
    if (sigma == 0.0) {
        return 1.0;
    }
    return (n/sigma) * exp(-0.5*squared((x-mu)/sigma));
}
float t = round(T * 100.0f);
auto* v = new float[dims] {0.0}; \\ the encoding vector
for (int i = 0; i < s; ++i){
    int currentPeak = (int) round(S[s] * 100.0f);
    int minPeak = currentPeak - t > 0 ?
        currentPeak - t :
        0;
    int maxPeak = currentPeak + t < dims ?
        currentPeak + t :
        dims - 1;
    for (int j = minPeak; j <= maxPeak; ++j) {
        float currentVal = v[j];
        float newVal = normpdf((float) j,
                               (float) currentPeak,
                               t / 3.0f);
        v[j] = std::max(currentVal, newVal);
    }
}
```

The peptide encoding vector is calculated as follows (here implemented in the C++ language):

```
const int dims = 500000;
auto* v = new float[dims] {0.0}; \\ the encoding vector
int index = 0;
for (int i = 0; i < p; ++i){
    index = (int) round(P[p] * 100.0f);
    v[index] = 1.0f;
}
```

Furthermore, a graphical explanation of the mass spectrum encoding is given in Supplementary Figure 8 and the peptide encoding is shown in Supplementary Figure 9. The complete source code for encoding and matrix multiplication is given in https://github.com/hgb-bin-proteomics/CandidateSearch and https://github.com/hgb-bin-proteomics/CandidateVectorSearch.

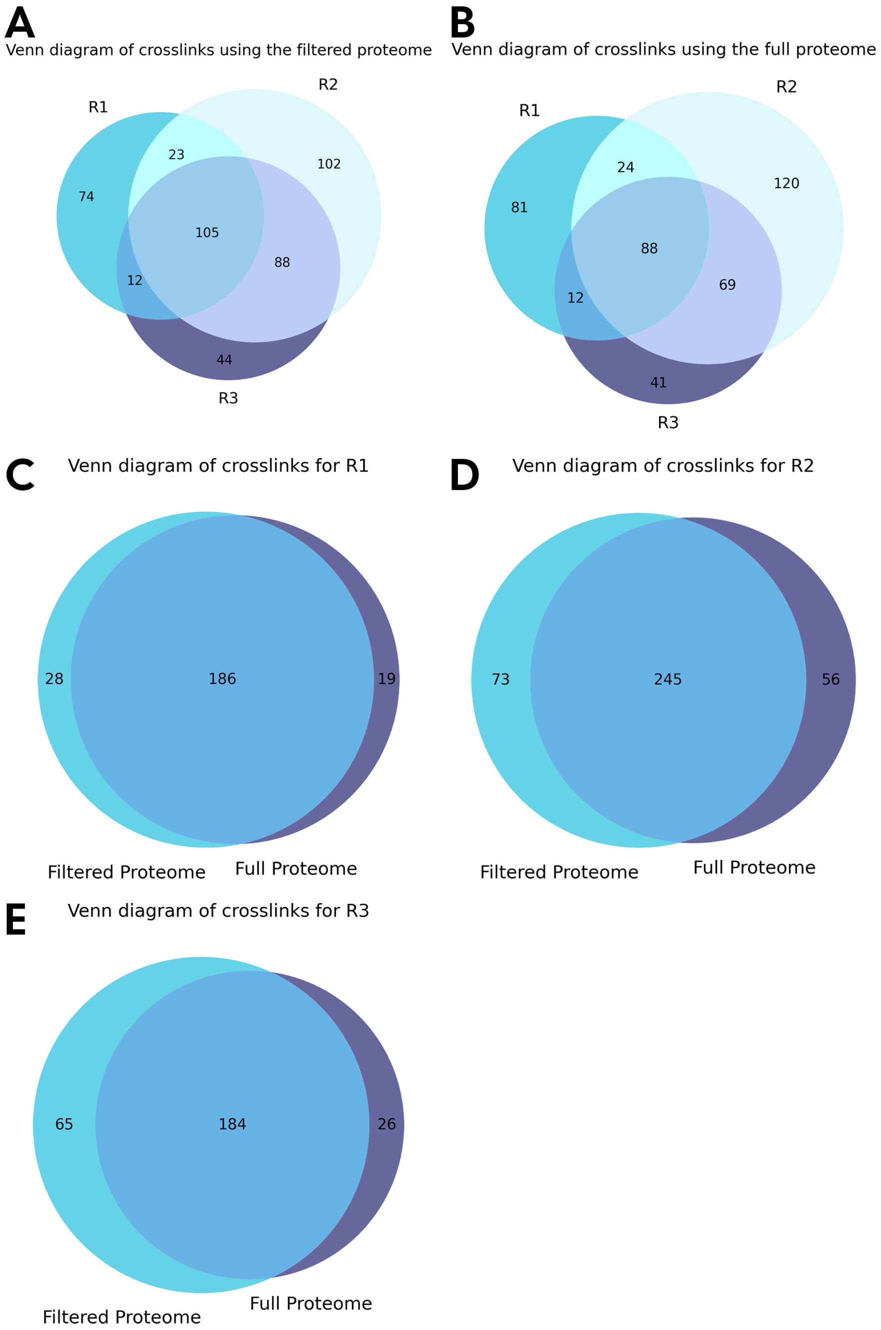


**Fig. 1** Overlaps of identified crosslinks at 1% FDR in the *C. elegans* dataset using MS Annika for search and xiFDR [1] for validation. A) Overlaps of identified crosslinks of the three biological replicates using the filtered proteome. B) Overlaps of identified crosslinks of the three biological replicates using the full proteome. C) Overlap of identified crosslinks for filtered and full proteome in replicate 1. D) Overlap of identified crosslinks for filtered and full proteome in replicate 2. E) Overlap of identified crosslinks for filtered and full proteome in replicate 3.

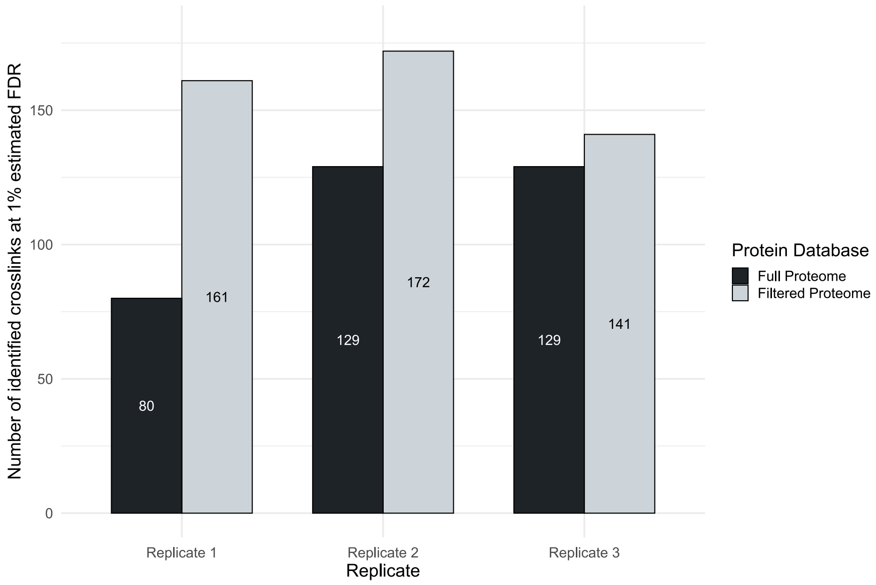


**Fig. 2** Influence of protein database size on the non-cleavable search of MS Annika when using the built-in valdiation algorithm. Mass spectrometry data of *C. elegans* nuclei was searched in MS Annika once using the full *C. Elegans* proteome (n = 26 695) and once using a filtered proteome of abundant proteins identified in a linear search (n = 3069). Results were validated for 1% estimated FDR using the built-in validation algorithm as described in [5]. The number of identified crosslinks varies a lot more compared to validation with xiFDR [1]. The biggest difference is observed for replicate one where the filtered database search reports 81 more crosslinks, more than double the amount of the search with the proteome-wide database. Interestingly, the difference is comparably small for replicate three, where the filtered proteome search only amounts in 12 more crosslink identifications. Overall 273 unique crosslinks are identified across the three replicates using the filtered proteome and 201 unique crosslinks using the full proteome.

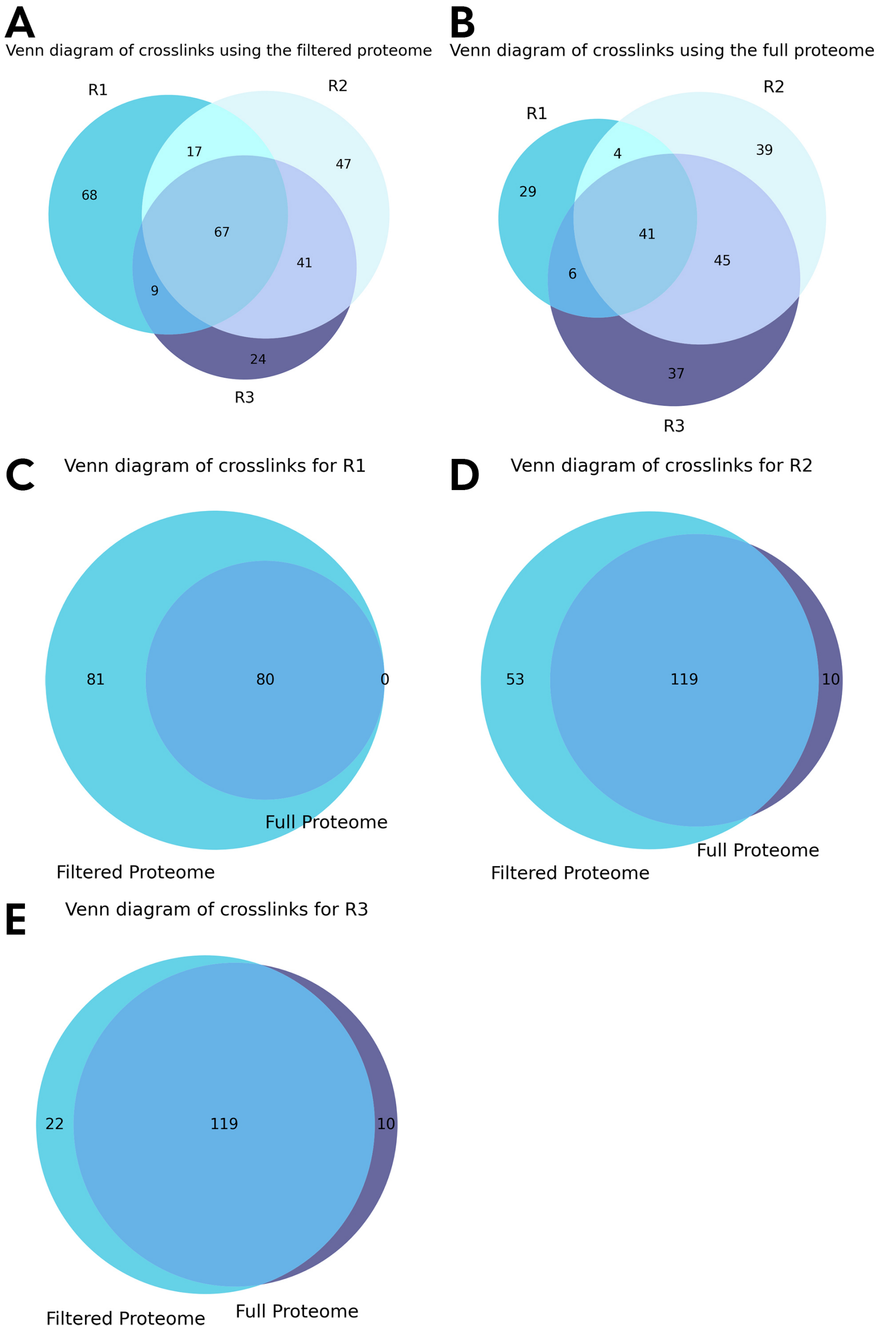


**Fig. 3** Overlaps of identified crosslinks at 1% FDR in the *C. elegans* dataset using MS Annika for search and validation. A) Overlaps of identified crosslinks of the three biological replicates using the filtered proteome. B) Overlaps of identified crosslinks of the three biological replicates using the full proteome. C) Overlap of identified crosslinks for filtered and full proteome in replicate 1. D) Overlap of identified crosslinks for filtered and full proteome in replicate 2. E) Overlap of identified crosslinks for filtered and full proteome in replicate 3.

**Table 1** Search times for crosslink identification in the three replicates of our *C. elegans* dataset using the full proteome-wide search.

| Replicate | Nr. of Mass Spectra | Search Time |
|---|---|---|
| Replicate 1 | 556013 | 5h 51min |
| Replicate 2 | 576443 | 6h 28min |
| Replicate 3 | 515924 | 5h 40min |

**Table 2** Hardware configuration of the system that we used for benchmarking and for the more demanding proteome-wide searches.

| Component | Product |
|---|---|
| MB | ASUS ROG Strix B650E-I |
| CPU | AMD Ryzen R9 7900X [12 cores @ 4.7 GHz base / 5.6 GHz boost] |
| RAM | Kingston 64 GB DDR5 RAM [5600 MT/s, 36 CAS] |
| GPU | ASUS Dual [Nvidia] GeForce RTX 4060 Ti OC [16 GB VRAM] |
| SSD/HDD | Corsair MP600 Pro NH 2 TB NVMe SSD [PCIe 4.0] |
| OS | Windows 11 Pro 64-bit (Version 10.0, Build 22631) |

**Table 3** Number of mass spectra containing diagnostic ions in the dataset by Beveridge and co-workers [2].

| Replicate | Nr. of Mass Spectra | Nr. of Mass Spectra with Diagnostic Ions |
|---|---|---|
| Replicate 1 | 5198 | 883 |
| Replicate 2 | 10538 | 2280 |
| Replicate 3 | 9892 | 1966 |

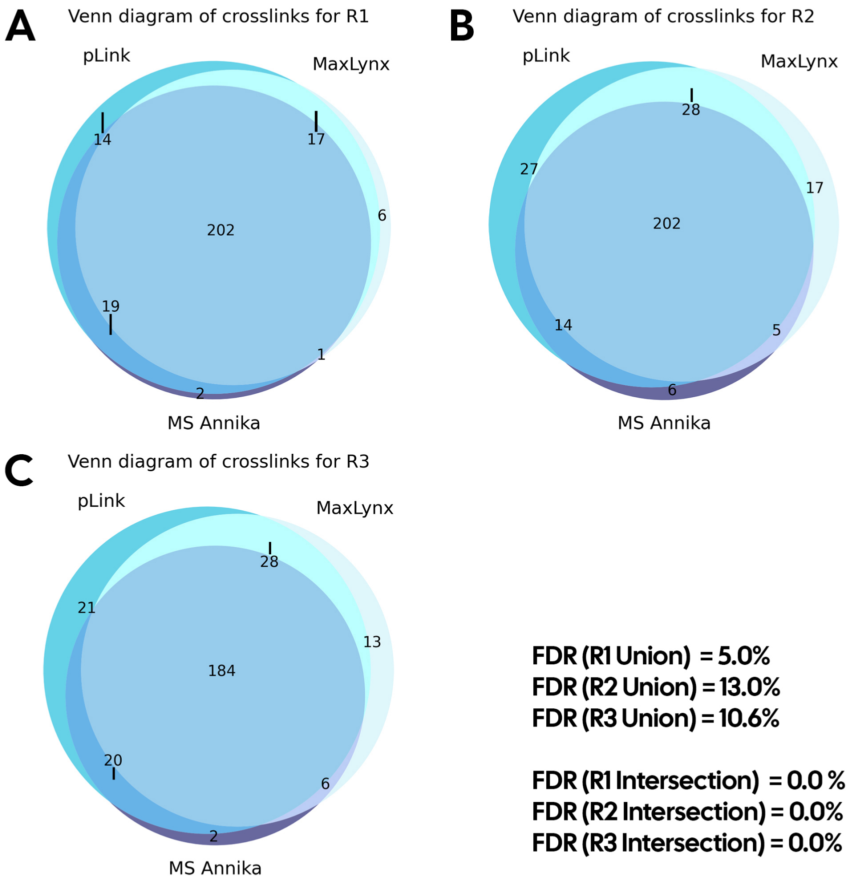


**Fig. 4** Overlaps of identified crosslinks at 1% FDR in the dataset by Beveridge and co-workers [2]. All venn diagrams show overlaps of identifications from the search engines MaxLynx [6], pLink [7] and MS Annika. A) Overlaps of crosslink identifications for replicate 1. B) Overlaps of crosslink identifications for replicate 2. C) Overlaps of crosslink identifications for replicate 3. The bottom right lists experimentally validated FDRs for unions and intersections: as expected the unions show higher experimentally validated FDRs while intersections report zero false positive hits.

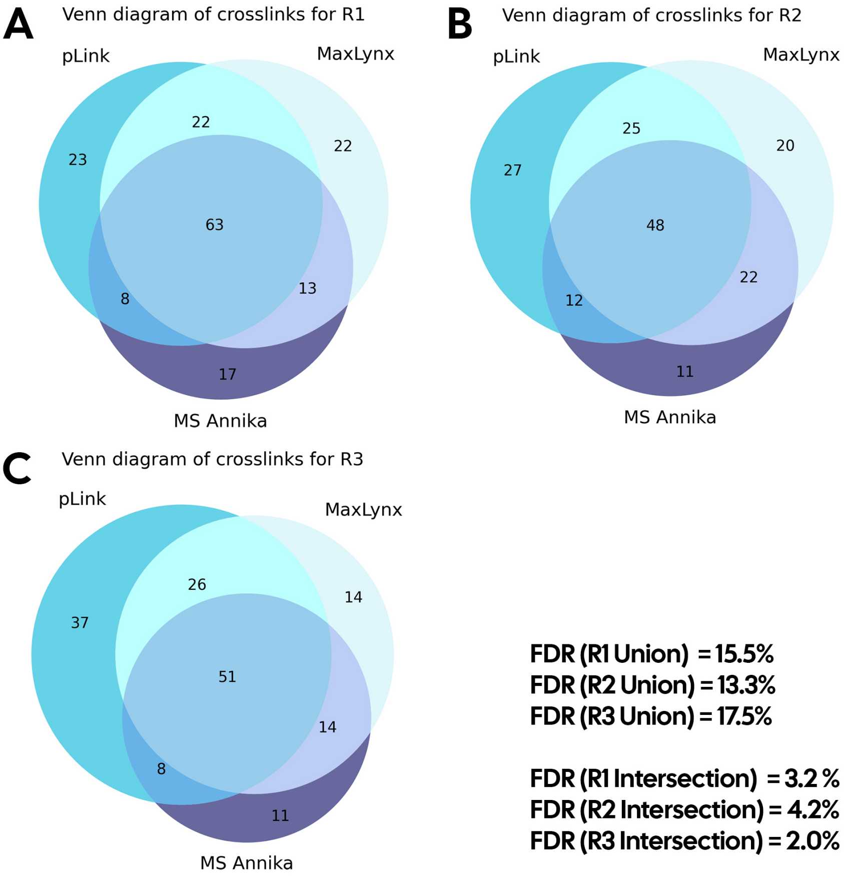


**Fig. 5** Overlaps of identified crosslinks at 1% FDR in the dataset by Matzinger and co-workers [3]. All venn diagrams show overlaps of identifications from the search engines MaxLynx [6], pLink [7] and MS Annika. A) Overlaps of crosslink identifications for replicate 1. B) Overlaps of crosslink identifications for replicate 2. C) Overlaps of crosslink identifications for replicate 3. The bottom right lists experimentally validated FDRs for unions and intersections: remarkably intersections still consist of up to two false positive hits and do not meet the 1% FDR criteria.

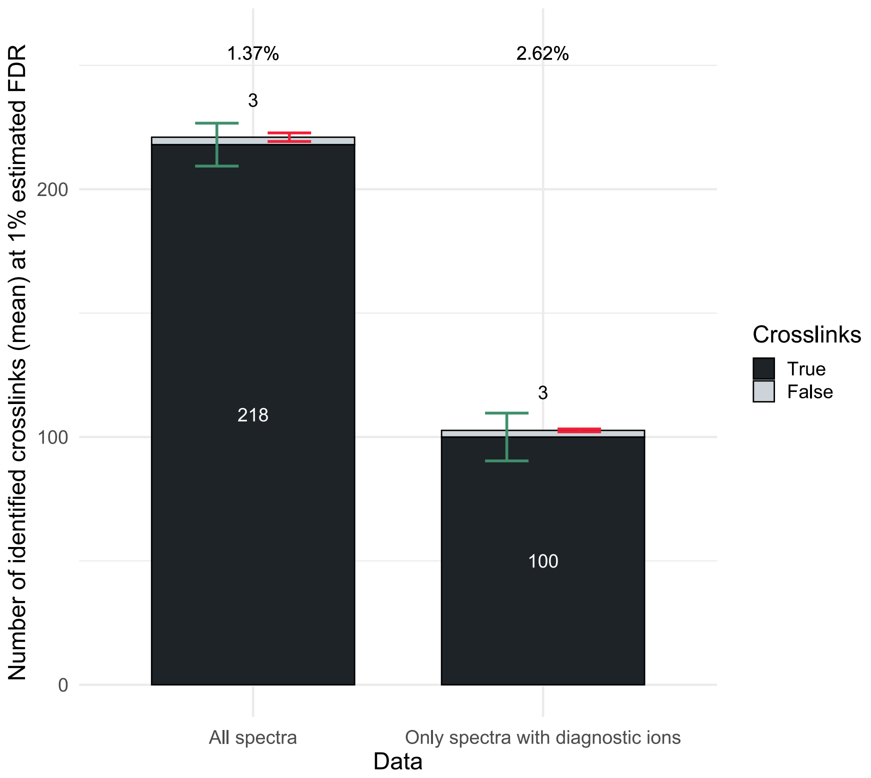


**Fig. 6** Comparison of running the MS Annika non-cleavable search on either all spectra or only spectra containing diagnostic ions of the benchmark dataset by Beveridge and co-workers [2]. Excluding spectra without diagnostic ions results in a substantial loss of identified true positive crosslinks, yielding less than half of the crosslinks that are detected when all spectra are searched. Because the number of false positive hits remains the same, this also constitutes a higher experimentally validated FDR of 2.62% and therefore an overall worse result. Results were validated for 1% estimated FDR. All numbers are averages from three technical replicates (n = 3), the number of crosslinks was rounded to the closest integer value. Percentage numbers above the bars denote the average calculated experimentally validated FDR rounded to two decimal places. Error bars denote the standard deviation, in green for true positive crosslinks and in red for false positive crosslinks.

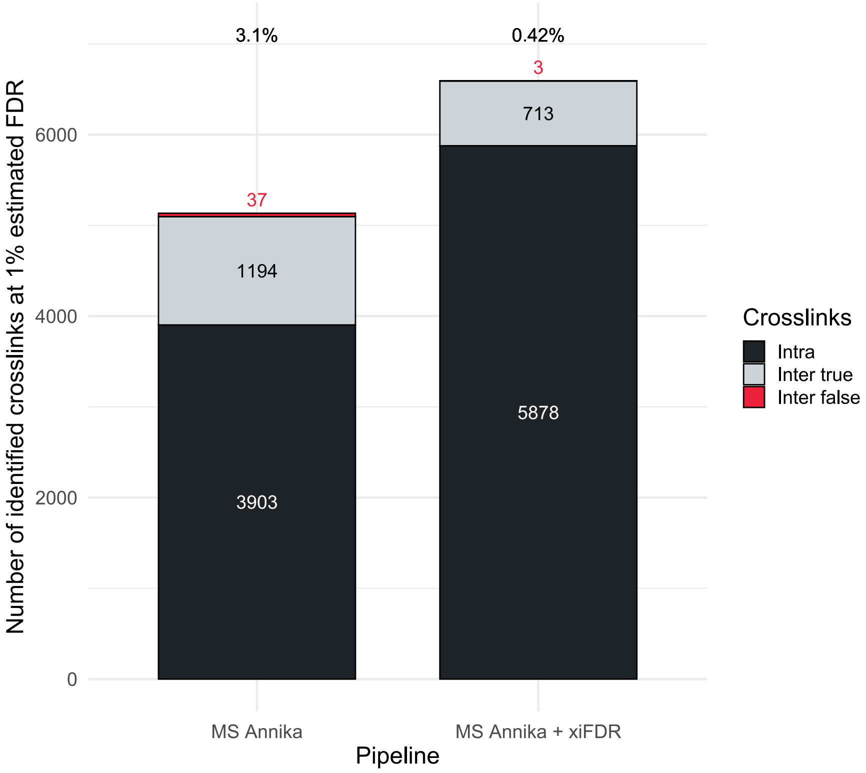


**Fig. 7** Comparison of crosslink results of MS Annika using either the built-in FDR validation or xiFDR [1] for validation, as demonstrated on the dataset by Lenz and co-workers [8]. Using more sophisticated validation approaches like xiFDR reduces the experimentally validated inter crosslink FDR from 3.1% to 0.42% while overall boosting the number of crosslink identifications from 5134 to 6594. Results were validated for 1% estimated FDR. Percentage numbers above the bars denote the calculated experimentally validated inter crosslink FDR rounded to two decimal places

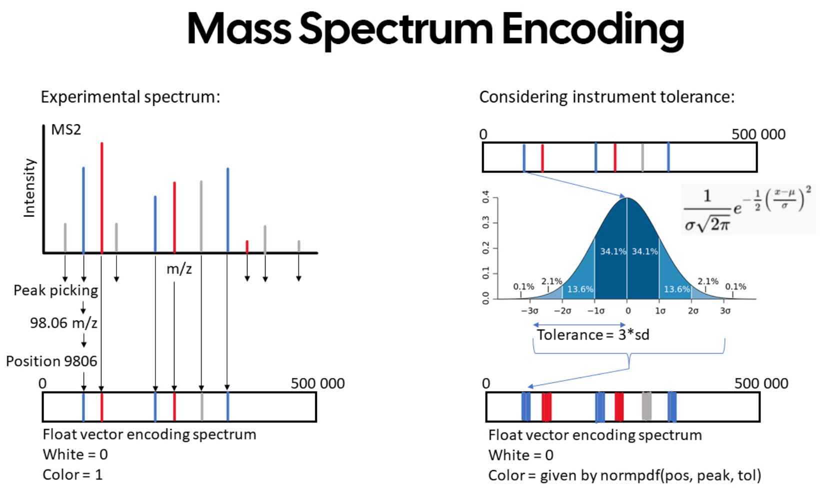


**Fig. 8** Visualization of the mass spectrum encoding. For every mass spectrum a 500 000-dimensional float vector is created and all values are initialized to zero. Mass spectra may be peak picked or otherwise preprocessed before encoding to avoid the incorporation of noise, e.g. by using the IMP MS2 Spectrum Processor node or other deconvolution algorithms. Subsequently, for every peak the m/z is taken, multiplied by 100 and rounded to the closest integer. If the resulting value is smaller than 500 000, the element at the vector index equal to the value is set to one. To account for instrument tolerance every peak is modelled as normal distribution with $\mu$ equal to the peak's m/z and $\sigma$ equal to tolerance/3, where tolerance is a user-definable parameter. The encoding works the same way, however instead of setting the elements in the vector to one, the resulting value of the normal distribution is taken, as shown on the right-hand side of the figure. Depiction of the normal distribution is taken from Wikimedia Commons, author: Ainali, licensed unter CC-BY-SA 3.0 https://commons.wikimedia.org/wiki/File:Standard_deviation_diagram_micro.svg.

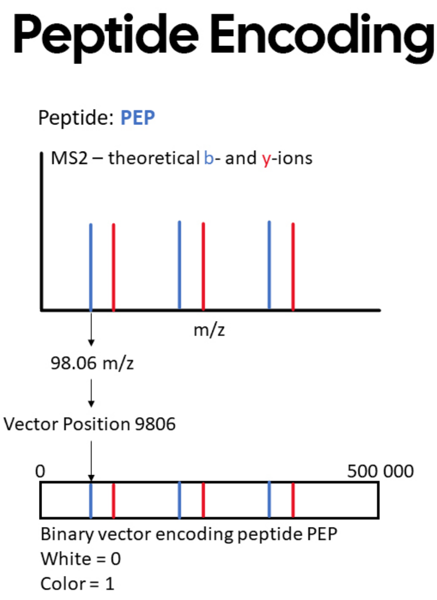


**Fig. 9** Visualization of the peptide encoding. Analogous to the mass spectrum encoding, every peptide is encoded as a 500 000-dimensional float vector. For every peptide such a vector is created and all values are initialized to zero. Furthermore, after calculating all theoretical ions for the given peptide, every ion's m/z value is multiplied by 100 and rounded to the closest integer. If the resulting value is smaller than 500 000, the element at the vector's corresponding index is set to one.

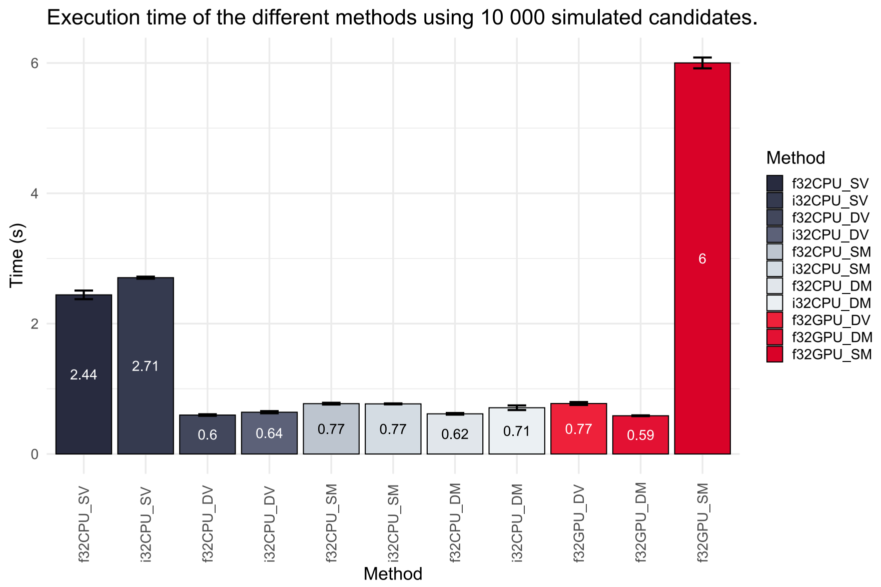


**Fig. 10** Synthetic benchmark of the different sparse matrix multiplication algorithms implemented in MS Annika. The benchmark consisted of simulating 10 000 peptides with 100 theoretical ions each, as well as 1001 simulated mass spectra with 1000 peaks each. For this benchmark the f32-based sparse matrix * dense matrix multiplication algorithm on the GPU performed best at an average of 0.59 seconds. The specific algorithms are described in the methods section in the main text. Values are averages of five different runs, error bars denote the standard deviation (n = 5). More synthetic benchmarks are available at https://github.com/hgb-bin-proteomics/CandidateVectorSearch/blob/master/benchmarks.md.

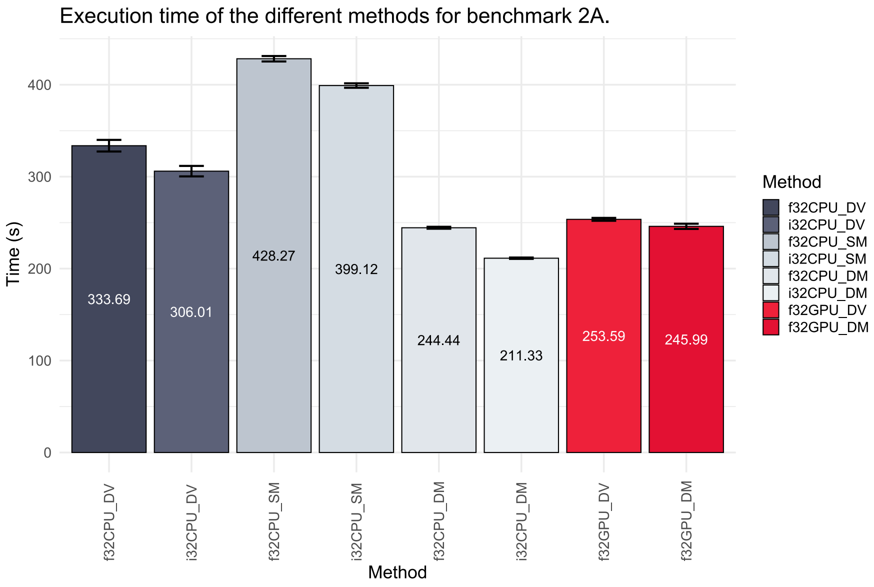


**Fig. 11** Benchmark of the different sparse matrix multiplication algorithms implemented in MS Annika using real data. Mass spectra of replicate 1 of the dataset by Beveridge and co-workers [2] were searched against a protein database consisting of the sequence of *S. pyogenes* Cas9 and the human SwissProt proteome. Some algorithms were omitted from benchmarking due to their long runtimes which were already immanent from the synthetic benchmarks, specifically *f32CPU_SV*, *i32CPU_SV* and *f32GPU_SM*. The results show that integer-based algorithms generally outperform the float-based counterparts and matrix * matrix multiplication outperforms matrix * vector multiplication. For our system (see 2) i32-based sparse matrix * dense matrix multiplication performed the best at an average of 211.33 seconds. The specific algorithms are described in the methods section in the main text. Values are averages of five different runs, error bars denote the standard deviation (n = 5). More benchmarks are available at https://github.com/hgb-bin-proteomics/CandidateSearch/blob/master/benchmarks.md.

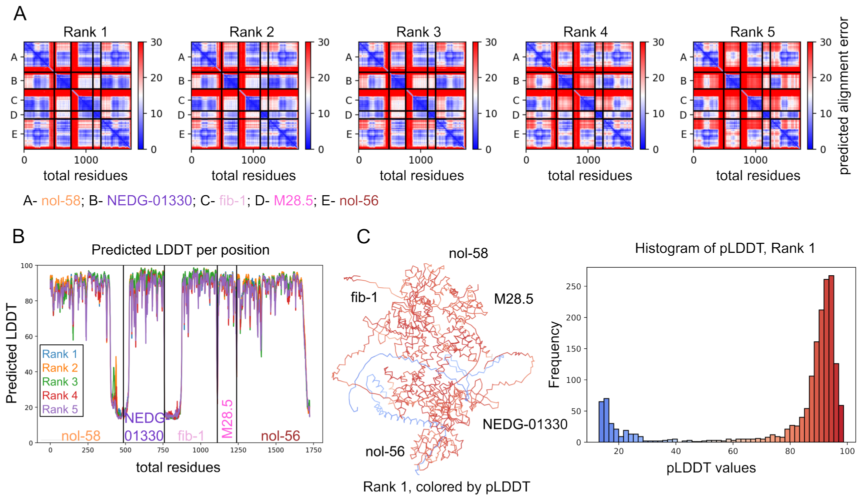


**Fig. 12** Diagnostic plots for the Box C/D RNP complex prediction. The nol-58, nol-56, M28.5 and fib-1 are predicted with high confidence to bind distinct binding sites. The interface between NEDG-01330 and nol-56 is also predicted with high confidence, while the n-terminal region of nol-56 and nol-58 resulted in low prediction accuracy. Shown are A: the PAE plot (Rank 1-5), B: the pLDDT plot of each rank separately, C: the structure of the top-ranked model in Ca trace coloured by pLDDT, and D: A histogram plot showing the distribution of pLDDT values of the Rank 1 structure, coloured by the confidence of the prediction (red: high confidence, blue: low confidence).

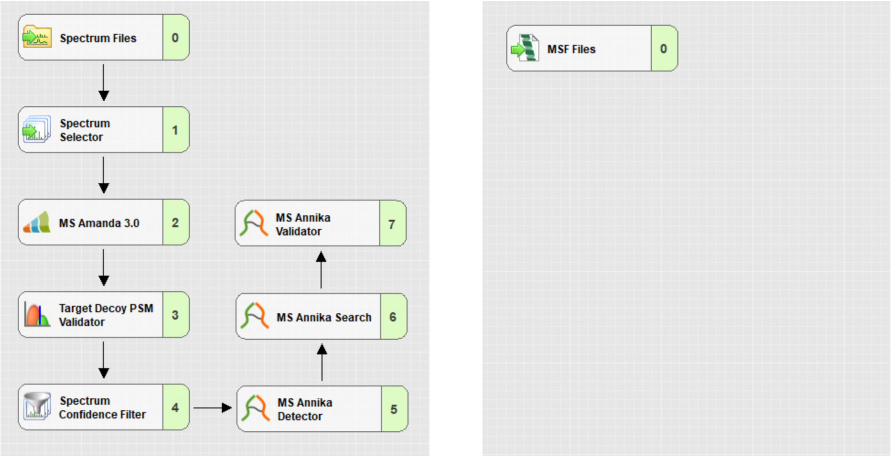


**Fig. 13** The standard Proteome Discoverer workflow for crosslink identification with MS Annika. Linear and monolinked peptides are first identified with MS Amanda [9, 10]. Mass spectra with associated high-confidence peptide spectrum matches (PSMs) are filtered out and not considered for crosslink search. Crosslinks are identified and validated with the MS Annika Detector node, the MS Annika Search node, and the MS Annika Validator node. The left panel shows the processing workflow while the right panel shows the consensus workflow.

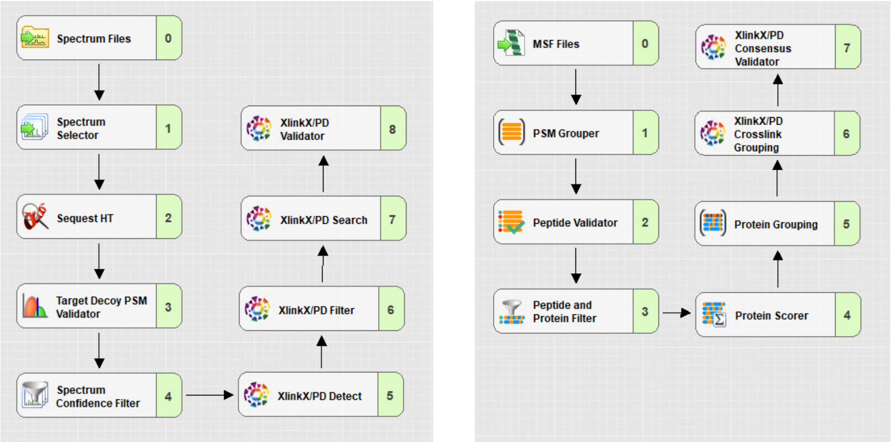


**Fig. 14** The non-cleavable HCD/CID MS2 Proteome Discoverer workflow for crosslink identification with XlinkX [4]. Linear and monolinked peptides are first identified with the Sequest HT node [11]. Mass spectra with associated high-confidence peptide spectrum matches (PSMs) are filtered out and not considered for crosslink search. Crosslinks are identified and validated with the XlinkX nodes. The left panel shows the processing workflow while the right panel shows the consensus workflow.

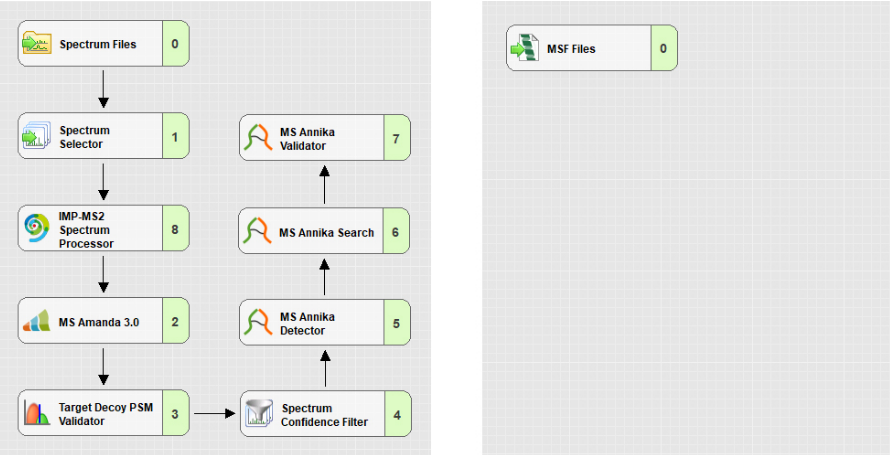


**Fig. 15** The Proteome Discoverer workflow for crosslink identification with MS Annika for complex samples. Mass spectra are first deisotoped with the IMP MS2 Spectrum Processor node. Then linear and monolinked peptides are identified with MS Amanda [9, 10] and mass spectra with associated high-confidence peptide spectrum matches (PSMs) are filtered out and not considered for crosslink search. Crosslinks are identified and validated with the MS Annika Detector node, the MS Annika Search node, and the MS Annika Validator node. The left panel shows the processing workflow while the right panel shows the consensus workflow.
